# Aberrant Fibro-Adipogenic Progenitor Subpopulations Drive Volumetric Muscle Loss-Induced Fibrosis

**DOI:** 10.1101/2025.05.11.653339

**Authors:** Shannon E. Anderson, Lauren A. Hymel, Hongmanlin Zhang, Jay M. McKinney, Thomas C. Turner, Mahir Mohiuddin, Woojin M. Han, Nan Hee Lee, Jeongmoon J. Choi, Gunjae Jeong, Emily Greenwood, Paramita Chatterjee, Seul Lee, Greg Gibson, Levi B. Wood, Edward A. Botchwey, Young C. Jang, Nick J. Willett

**Affiliations:** Wallace H. Coulter Department of Biomedical Engineering, Georgia Institute of Technology and Emory University, Atlanta, GA, USA 30332; Parker H. Petit Institute for Bioengineering and Bioscience, Georgia Institute of Technology, Atlanta, GA, USA 30332; Atlanta Veterans Affairs Medical Center, Decatur, GA, 30030; Department of Orthopaedics, Emory University, Atlanta, GA, USA 30322; School of Materials Science and Engineering, Georgia Institute of Technology, Atlanta, GA, USA 30332; School of Biological Sciences, Georgia Institute of Technology, Atlanta, GA, USA 30332; Department of Orthopaedics, Icahn School of Medicine at Mount Sinai, New York, NY, USA 10029; Black Family Stem Cell Institute, Icahn School of Medicine at Mount Sinai, New York, NY, USA 10029; Center for Integrative Genomics, School of Biological Sciences, Georgia Institute of Technology, Atlanta, GA, USA 30332; Marcus Center for Therapeutic Cell Characterization and Manufacturing, Georgia Institute of Technology, Atlanta, GA, USA 30332; George W. Woodruff School of Mechanical Engineering, Georgia Institute of Technology, Atlanta, GA, USA 30332; Phil and Penny Knight Campus for Accelerating Scientific Impact, University of Oregon, Eugene, OR, USA 97403; The Veterans Affairs Portland Health Care System, Portland, OR, USA 97239; Emory University School of Medicine, Atlanta, GA, USA 30322

## Abstract

Volumetric muscle loss (VML) injuries result in chronic fibrosis, inflammation, and persistent functional deficits. Fibro-adipogenic progenitor (FAP) cells are a heterogeneous, muscle-resident stromal cell population that play a crucial role in muscle regeneration, but also contribute to fibrosis in muscle disease. The role of FAPs in VML is not well established and may be critical target to ensure functional muscle regeneration after VML. We utilized a VML model in the mouse quadriceps to study the location, secretome, surface marker distribution, gene expression, and single-cell transcriptional profile of FAPs after VML. After VML, a subpopulation of FAPs highly expressed β1-integrin and were elevated in the post-VML muscle tissue; these FAPs had increased fibrotic gene expression and increased myofibroblast differentiation potential. Transforming growth factor-β1 (TGF-β1) and tissue inhibitor of matrix metalloproteinase 1 (TIMP1) were identified as secreted proteins from VML derived FAPs that produced both pro-fibrotic and anti-myogenic signaling. These data establish an aberrant FAP sub-population that are elevated in VML injury and provides novel targets for future scarless muscle regeneration in VML.

## Introduction

The interaction between muscle stem cells (MuSCs) and FAPs is critical to the regenerative process following skeletal muscle injury (*1*). FAPs are a muscle-resident population of mesenchymal cells that can differentiate into fibroblasts or adipocytes; during homeostasis they reside as non-committed progenitor cells in the space between myofibers (*2*). FAPs function as critical regulators of skeletal muscle during homeostasis and regeneration by secreting various pro-myogenic factors, including Wnt1-inducible signaling protein 1 (WISP1) and follistatin (*3, 4*). The ablation of FAPs from the muscle results in significant skeletal muscle atrophy (*2*). Following muscle injury, FAPs typically proliferate, peaking around 4 days post injury, and create a transitional niche for MuSCs before returning to pre-injury levels by 14 days post-injury (*1, 5*). However, during chronic injuries or aging, FAPs have been shown to persist, differentiate into myofibroblasts and adipocytes, and contribute to pathological fibrosis and fatty infiltration (*1, 6, 7*).

FAPs persistence, function, and differentiation is closely related to the inflammatory state of the tissue. Pro-inflammatory macrophage secretion of tumor necrosis factor α (TNF-α) is required for efficient FAP clearance by apoptosis; however, the persistent presence of both TNF-α and transforming growth factor beta 1 (TGF-β1) leads to FAPs survival and accumulation (*8*). Environments containing both pro- and anti-inflammatory cytokines are characteristic of chronic inflammation and dysregulated macrophage polarization (*8, 9*) – a state that is believed to be similar to the dysregulated immune environment in VML (*10*). FAPs have also been identified as a heterogeneous cell population with different subpopulations playing a role in the progression of disease states. In a mouse model of muscular dystrophy, a vascular cell adhesion molecule 1 (Vcam1)-expressing, pro-fibrotic subtype of FAPs was previously identified to accumulate following a disruption in the immune response (*11*). While FAPs contribute to fibrosis after minor acute injury, the ablation of FAPs or fibroblasts impairs muscle regeneration (*2, 12*). This previous literature outlines dueling roles for FAPs as either a pro-myogenic or pro-fibrotic population following muscle injury.

The accumulation of fibrotic tissue, persistent inflammation (*13–15*), and chronic upregulation of TNF-α (*14*) and TGF-β1 are all hallmarks of VML injuries (*16*). Previously, we established a critical threshold size for VML defects in the mouse quadriceps; defects larger than the threshold led to VML injuries with persistent inflammation, fibrosis, and a lack of re-innervation (*15*). The persistent inflammation and eventual fibrosis and fatty infiltration in VML injury implicate a dysregulated state of FAPs; however, this has not been well characterized. Recently, it was reported that FAPs following VML increase in number and that those FAPs have increased gene expression related to proliferation and fibrosis but no significant differences in pro-myogenic cytokines insulin-like growth factor-1 (IGF-1) or follistatin (*17*). As FAPs have previously been found to be a heterogeneous population in skeletal muscle, it is likely that the promotion of myogenesis over fibrosis requires the balancing of FAPs subpopulations. Thus, it is crucial to probe the inherent FAP heterogeneity induced within highly inflammatory microenvironments to uncover their subset-specific roles in muscle regeneration.

Herein, we characterize the temporal systemic and local response of MuSCs, FAPs, and related cytokines following critical VML. We then sought to investigate the transcriptional, protein, and secretory heterogeneity of FAPs following VML to identify a subpopulation which may contribute to fibrosis and impair myogenesis. We hypothesized that FAPs from critical VML injury would present with an aberrant pro-fibrotic phenotype pre-disposed towards fibrosis.

## Results

### VML injury results in elevated blood serum cytokines, activated MuSCs, and persistent FAPs

Flow cytometry analyses were conducted on mice following a recoverable, subcritical injury (2mm biopsy) or irrecoverable VML defect (3mm biopsy) to the quadriceps at 1, 3, or 7 days post-injury (n=4 per group, Fig. 1A). Biplot gating of analyzed events was done to determine the number of MuSCs and FAPs present per milligram of muscle tissue, or cell concentration within the injured muscles. At 7 days post-injury, MuSC concentration was significantly elevated in critically-sized VML injured quadriceps (Fig. 1B, *p*<0.01). FAPs in subcritical and critical injuries were present at relatively low levels at 1-day post-injury (Fig. 1C). By 3 days post-injury, the number of FAPs in critically sized injuries increased to a level significantly greater than in subcritical injures (*p*<0.001). Four days later, subcritically injured muscle had no significant difference in FAPs concentration to muscle at 1-day post-injury, whereas muscle from critically-sized VML injuries maintained a persistent significant elevation in FAPs levels (*p*<0.001).

**Fig. 1.**
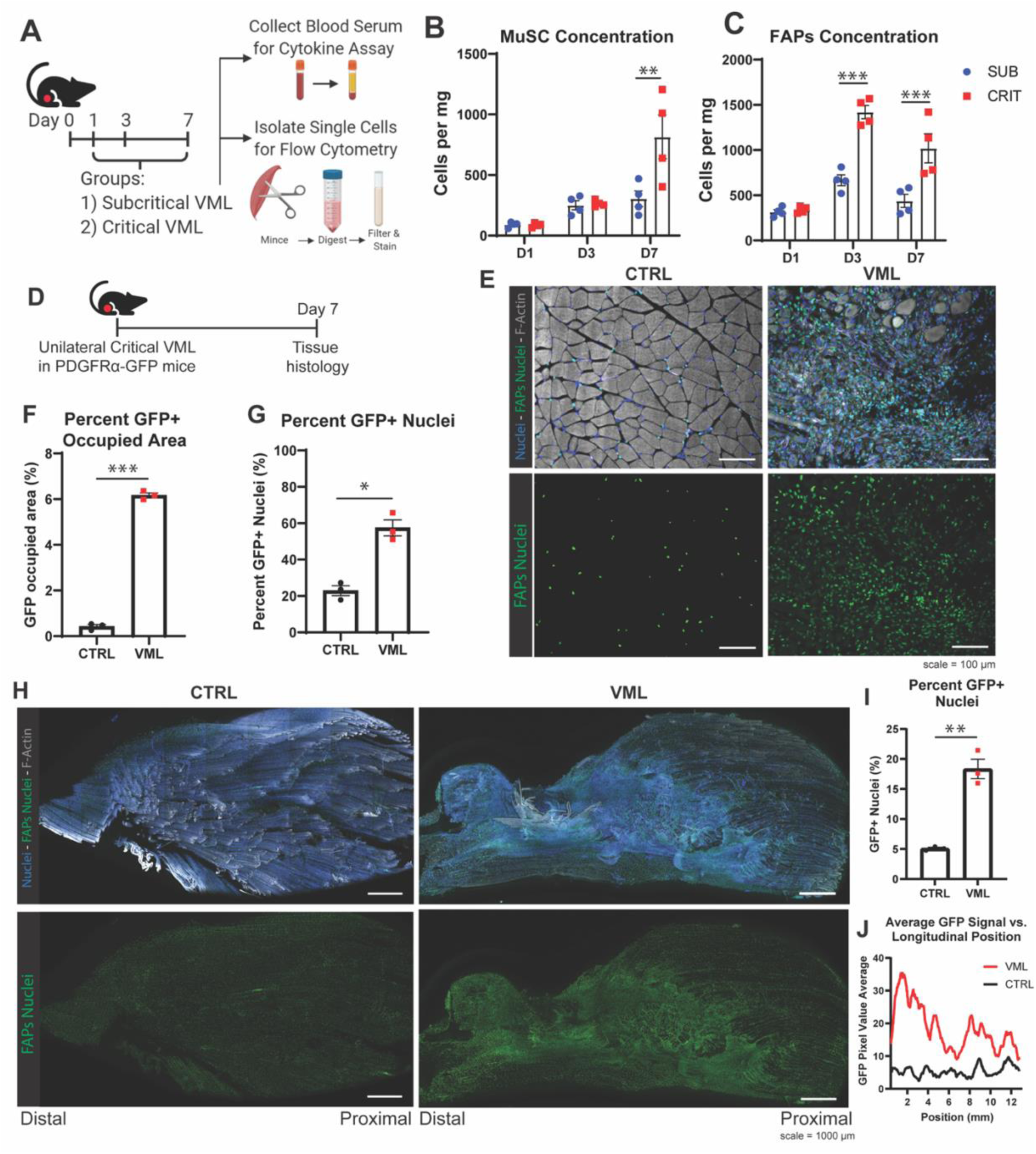
FAPs are significantly elevated following VML injury and are pervasive throughout the muscle. Experimental timeline (**A**) for flow cytometry and cytokine analysis following injury at indicated timepoints. Biplot gating was used to quantify MuSCs (**B**) and FAPs (**C**) over time following subcritical (SUB) or critical (CRIT) injury. Cells and serum were from day 1, day 3, or day 7 (D1, D3, D7 respectively) following injury. Experimental timeline (**D**) of PDGFRα-GFP unilateral critical VML with tissue analysis at 7 days post-injury. Representative cross-section images (**E**) of contralateral control (CTRL) and critical VML (VML) injured muscle. Quantification of cross section images as percent GFP occupied area (**F**) and percent GFP^+^ nuclei (**G**). Representative longitudinal images (**H**) of contralateral (CTRL) and critical VML injured (VML) tissue. Quantification of GFP^+^ FAPs in longitudinal sections as percent GFP^+^ nuclei (**I**) and average pixel value versus position (**J**, distal to proximal, 0-13 mm). IHC staining for nuclei (blue), f-actin (grey), and FAPs PDGFRα-GFP nuclei (green). Statistics in GraphPad Prism 8, students t-test (**F, G** and **I**), two-way ANOVAs (**B** and **C**) were performed on log transformed raw data to correct for inequality in variance using Sidak post-hoc test for multiple comparisons. *=*p*<0.05, **=*p*<0.01, ***=*p*<0.001. n=3 or 4 animals per experimental group.

The dimensionality reduction technique, spanning-tree progression analysis of density-normalized events (SPADE), was used to cluster MuSC flow cytometry data at all time points to determine changes in the MuSC population after injury. The SPADE tree showed a delineation between forward scatter area (FSC-A) high and FSC-A low MuSCs, which indicates a difference in cell size (Fig. S1A). The events at the FSC-A high nodes were then broken down by their respective experimental groups and compared. The percentage of total MuSCs which were FSC-A high (cells which were larger in size), increased over time from 1-day post-injury to 3 days post-injury similarly in both subcritical and critical injuries. However, from 3 to 7 days post-injury, the percentage of FSC-A high MuSCs remained constant in subcritical injuries but continued to increase in critically injured muscle. At 7 days post-injury, the percentage of FSC-A high MuSCs from muscle with a critically-sized injury was significantly greater than those in subcritical injuries (Fig. S1B, *p*<0.001). The percentage of FSC-A high MuSCs in critical injuries was significantly elevated above the percentage in uninjured controls at all time points, whereas the same was only true in subcritical injuries at 3 and 7 days post-injury (Fig. S1B, *p*<0.05). When MuSCs transition from a quiescent state to an activated state, the size of the cell increases (Fig. S1C) (*18*), suggesting that critical VML specifically induces an increased concentration of predominantly activated MuSCs compared to subcritical VML.

To further investigate the fusion capacity of MuSCs following subcritical and critical VML, we utilized a mouse model with tamoxifen inducible TdTomato expression in paired box transcription factor 7 (*Pax7*) positive MuSCs (*Pax7^CreERT^;R26^lslTdTomato^*). This mouse model allowed for the tracking of cells derived from an MuSC lineage at the time of the VML injury. 14 days following injury, tissue was used for whole mount, longitudinal imaging (Fig. S1D). Contralateral control muscle at both low and high magnification (Fig. S1, E and F) showed TdTomato^+^ MuSCs along myofibers, and some low expression of TdTomato^+^ myofibers, indicating MuSC fusion. Following subcritical injury, high expression of TdTomato^+^ myofibers can be clearly seen, even at lower magnification (Fig. S1, G and H). The highest expressing fibers were located in the middle of the quad, nearest to where the injury would have been induced (Fig. S1H, middle) indicating the highest levels of MuSC fusion within the defect. Conversely, further away from the injury, the fibers are less brightly red and more MuSCs can be seen along myofibers (Fig. S1H, distal and proximal). In VML, TdTomato^+^ myofibers are present at all regions along the length of the muscle but do not bridge the defect area (Fig. S1, I and J). High magnification images showed MuSCs are proliferating, activating, and fusing following both critical VML and subcritical injury, indicating that MuSCs are responding to the injury and attempting to regenerate muscle.

While critically-sized VML defects have a significantly elevated concentration of MuSCs (Fig. 1B), the majority of the MuSCs were adjacent to the defect site (Fig. S1). To further assess resident muscle cells in the injured tissue and in the defect region, we next analyzed FAPs. To evaluate the localization of FAPs within critically injured muscle, we utilized a reporter mouse which expresses enhanced green fluorescent protein (EGFP) within the nuclei of cells expressing platelet-derived growth factor receptor α (PDGFRα, referred to as PDGFRα^EGFP^ mice). Unilateral VML injuries were performed on PDGFRα^EGFP^ mice and the number and location of FAPs at 7 days post-injury throughout the tissue was imaged and quantified (Fig. 1D-J). In VML injured tissue, GFP^+^ signal occupied about 6% of the total image area, a significantly higher percentage than in contralateral control tissue (Fig. 1F, *p*<0.001). Additionally, the percentage of nuclei which were GFP^+^ was measured and compared between VML and contralateral control sections. In the VML injury area, approximately 60% of nuclei were GFP^+^, indicating they were FAPs or FAPs-derived cells. This was significantly higher than in contralateral control tissue, where approximately 20% of nuclei were GFP^+^ (Fig. 1G, *p*<0.05).

Longitudinal whole mount sections allowed for visualization of FAPs throughout the length of the quadriceps muscle in critically injured and contralateral control tissue (Fig. 1H). The percentage of GFP^+^ nuclei was quantified in these sections. Approximately 15% of nuclei along the length of critically injured muscle were GFP^+^, significantly higher than the 5% of GFP^+^ nuclei in contralateral control tissue (Fig. 1I, *p*<0.05). The GFP fluorescence intensity was also measured as average pixel value and plotted along the length of the muscle from distal (0mm) to proximal (13mm) (Fig. 1J). The increased fluorescence GFP intensity illustrates the higher concentration of FAPs distally and within the injury compared to tissue proximal to the defect.

To determine whether critical VML in our murine quadriceps model induces a marked systemic immune response, we measured the cytokine levels in blood serum following subcritical injury and critical VML using a Luminex multiplex ELISA assay. Twelve cytokines were present at detectable levels in blood serum at days 1, 3, and 7 post muscle injury (Fig. 2A). Partial least square discriminant analysis (PLSDA) was used to separate each injury size and time point along two latent variables (LVs) (Fig. 2B). Individual sample scores on LV1 and LV2 were grouped by injury size and timepoint and compared (Fig. 2, C and E). There were no significant differences in scores between groups on LV1 (Fig. 2C). On LV2, there were significant differences between each time point (days 1, 3, and 7) in both subcritical and VML injuries, with scores on LV2 increasing with time (Fig. 2E, *p*<0.05). At day 1 post-injury, subcritical injuries had significantly lower LV2 scores than critical injuries (Fig. 2E, *p*<0.05). Loading plots were generated to show the relative weighting of each cytokine used to separate samples along LV1 and LV2 (Fig. 2, D and F). Interleukin (IL)-1α, lipopolysaccharide-induced CXC chemokine (LIX, also known as CXCL5), and monokine induced by interferon gamma (MIG, also known as CXCL9) contributed to positive LV2 scores (Fig. 2F). Conversely, IL-6, granulocyte colony stimulating factor (G-CSF), keratinocyte chemoattractant (KC), interferon gamma-induced protein 10 (IP-10), TNFα, IL-5, IL-13, and IL-17 contributed to negative scores of LV2 (Fig. 2F).

**Fig. 2.**
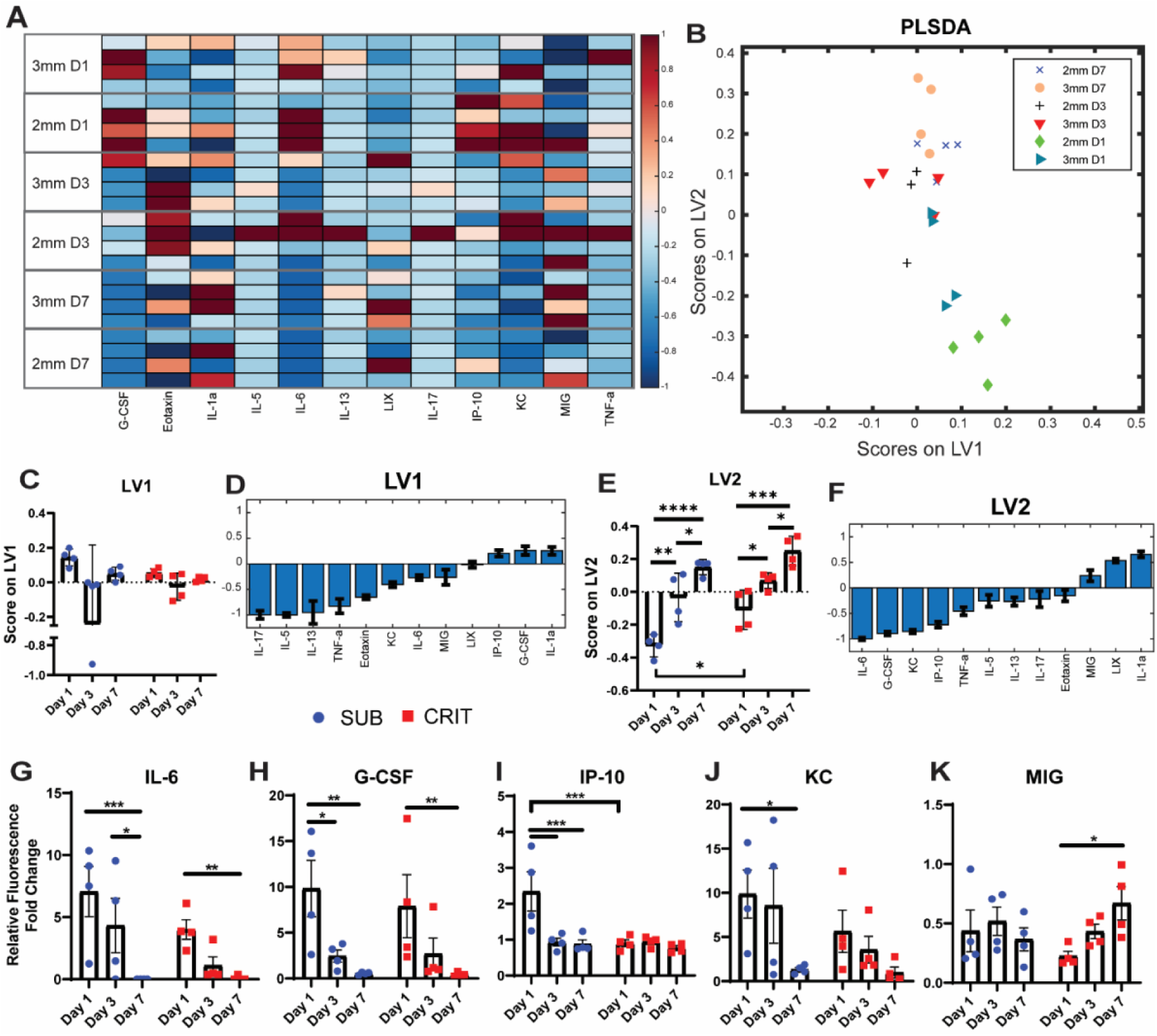
Blood serum cytokines fluctuate similarly over time following subcritical and critical VML. Luminex cytokines multiplex assay was used to measure the cytokine concentration in serum of mice following critical (CRIT, 3mm biopsy) and subcritical (SUB, 2mm biopsy) VML. Measured fluorescence was normalized to the level in uninjured control serum, values reported as fold change. All cytokine levels were plotted by their z score expression value in a heat map (**A**) with red color indicating higher expression and blue indicating lower expression. (**B**) PLSDA was used to reduce dimensionality and plot each sample based on their score on latent variables (LV) 1 and 2. Scores of each sample of LV1 and 2 are plotted (**C** and **E**) and the loading of each cytokine in each LV are shown (**D** and **F**). Fluorescence fold change to uninjured controls are compared between groups for significant differences and plotted for IL-6 (**G**), G-CSF (**H**), IP-10 (**I**), KC (**J**), and MIG (**K**). Two-way ANOVA performed in GraphPad Prism 8 with Sidak post-hoc test, *=*p*<0.05, **=*p*<0.01, ***=*p*<0.001, ****=*p*<0.0001. n=4 animals per experimental group.

Fold change in individual cytokine levels to muscle from uninjured control animals were compared over time and between groups via a two-way ANOVA. IL-6, G-CSF, and KC had similar cytokine expression patterns over time in subcritical and critical VML injuries, with significant differences only within injury sizes between time points (Fig. 2, G, H and J). IP-10 was expressed at significantly higher levels in subcritical injuries at 1-day post-injury than at later time points in the same injury size and critical VML injuries at the same time point (Fig. 2I, *p*<0.05). MIG was present at consistent levels following subcritical injury, but at levels which increased over time following critical VML injury reaching a concentration at 7 days post-injury which was significantly greater than 1-day post-injury (Fig. 2K, *p*<0.05).

### FAPs isolated from VML injured muscles have an altered secretome and increased differentiation potential in response to TGF-β1

As FAPs concentration is significantly increased in VML injured muscles (Fig. 1C), we investigated the secretory and differentiation behaviors of FAPs cultured alone. These functional differences were evaluated on FAPs sorted via fluorescence activated cell sorting (FACS) from uninjured mice or from VML injured PDGFRα^EGFP^ mice 7 days post-injury and then the isolated FAPs were subsequently cultured *in vitro* (Fig. 3A). Conditioned media was collected immediately before differentiation was induced after 6 days in culture and cytokine concentrations were measured. Cell culture media from wells with VML derived FAPs had significantly elevated concentrations of KC, VEGF, MCP-1, and TGF-β1 compared to media from wells with uninjured FAPs (Fig. 3B, *p*<0.05). Fibrotic differentiation was induced by a reduction in serum content in the media with or without the addition of TGF-β1. Immunohistochemical (IHC) staining following differentiation for myofibroblast marker alpha smooth muscle actin (αSMA) or adipocyte marker perilipin (Fig. 3C) showed that VML derived FAPs had a higher mean fluorescence expression of αSMA in the presence of TGF-β1 when compared to uninjured FAPs (Fig. 3D). However, without TGF-β1 stimulus, VML derived FAPs had increased perilipin expression, which was significantly reduced by TGF-β1 stimulus (Fig. 3E). The fraction of GFP^+^ nuclei was also quantified. GFP^+^ nuclei are indicative that FAPs are still in a progenitor phenotype expressing PDGFRα. Uninjured FAPs stimulated with TGF-β1 showed a significant decrease in the fraction of nuclei expressing GFP, but there was no such reduction in VML derived FAPs (Fig. 3F).

**Fig. 3.**
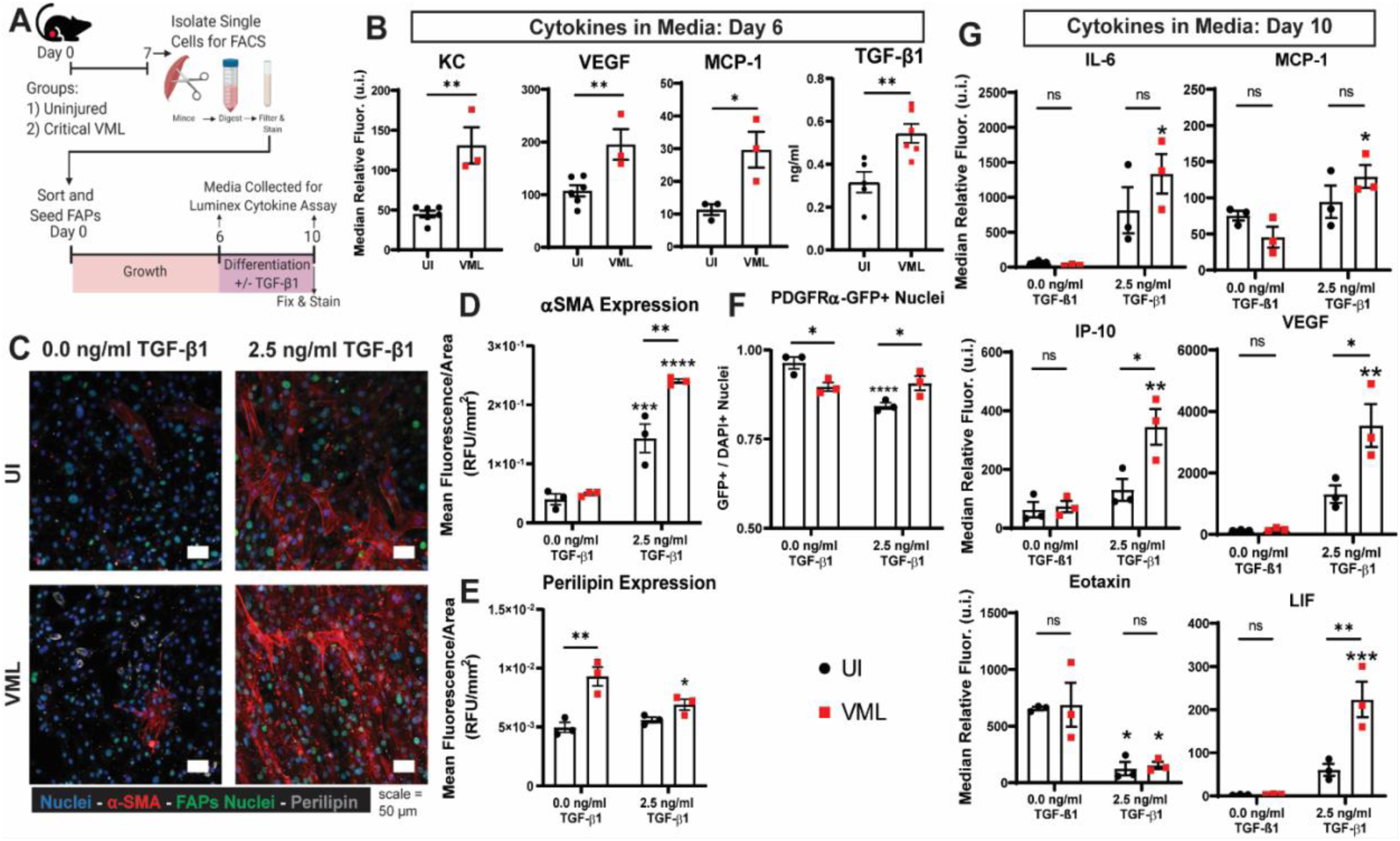
VML-derived FAPs have altered secretome and increased responsiveness to TGF-β1. Experimental design (**A**), briefly FAPs were isolated from uninjured (UI) or VML injured tissue 7 days post-injury. Isolated FAPs were then seeded and cultured for 10 days. Cytokine concentrations in conditioned media were measured at day 6 in culture (**B**) and cytokines with significant differences are shown. Representative IHC staining (**C**) for nuclei (blue), αSMA (red), FAPs nuclei (green), and perilipin (grey). Mean fluorescence intensity per area measured was quantified for αSMA (**D**) and perilipin (**E**) signal. The fraction of nuclei expressing GFP (**F**) was quantified. Cytokines in conditioned media at day 10, immediately before fixation, were measured (**G**) and cytokines with significant differences are shown. Statistics performed in GraphPad Prism 8, students t-test, *=*p*<0.05, **=*p*<0.01, ***=*p*<0.001, ****=*p*<0.0001. Asterisks above individual groups indicates a difference between that group’s unstimulated control, stars above comparison bars indicate differences between UI and VML in either stimulated or unstimulated condition. n=3-6 replicates per experimental group.

Following differentiation, conditioned media was collected for Luminex cytokine analysis. VML derived FAPs stimulated with TGF-β1 expressed significantly higher levels of IL-6, MCP-1, IP-10, VEGF, and leukemia inhibitory factor (LIF) when compared to VML derived FAPs not stimulated with TGF-β1 (Fig. 3G, *p*<0.05). By contrast, uninjured derived FAPs did not significantly increase their expression of any of these cytokines with TGF-β1 stimulus. In the case of IP-10, VEGF, and LIF, TGF-β1 stimulated VML derived FAPs had significantly increased expression over stimulated uninjured FAPs as well.

Injured VML muscle showed elevated MuSC levels within the injured tissue (Fig. 1B), increased MuSC fusion surrounding the defect (Fig. S1), and higher numbers of FAPs in areas of non-bridging following critical VML (Fig. 1E); given these observations we sought to characterize the cross-talk of VML derived FAPs and MuSCs using an *in vitro* model. MuSCs isolated from β-actin-GFP mice were cultured for three days and then allowed to differentiate either alone, cultured with FAPs in transwell from uninjured muscle, or FAPs in transwell from day 7 VML muscle for an additional five days (Fig. 4A). The number of multinucleated myotubes and total nuclei were quantified after eight days of MuSC culture (Fig. 4, B and C). MuSCs showed a significant increase in myotube and nuclei number when uninjured FAPs were cultured in transwell (Fig. 4, B and C, *p*<0.05), a phenomenon which has been previously reported. However, VML derived FAPs did not result in a significant increase in myotube or nuclei number when cultured in transwell.

**Fig. 4.**
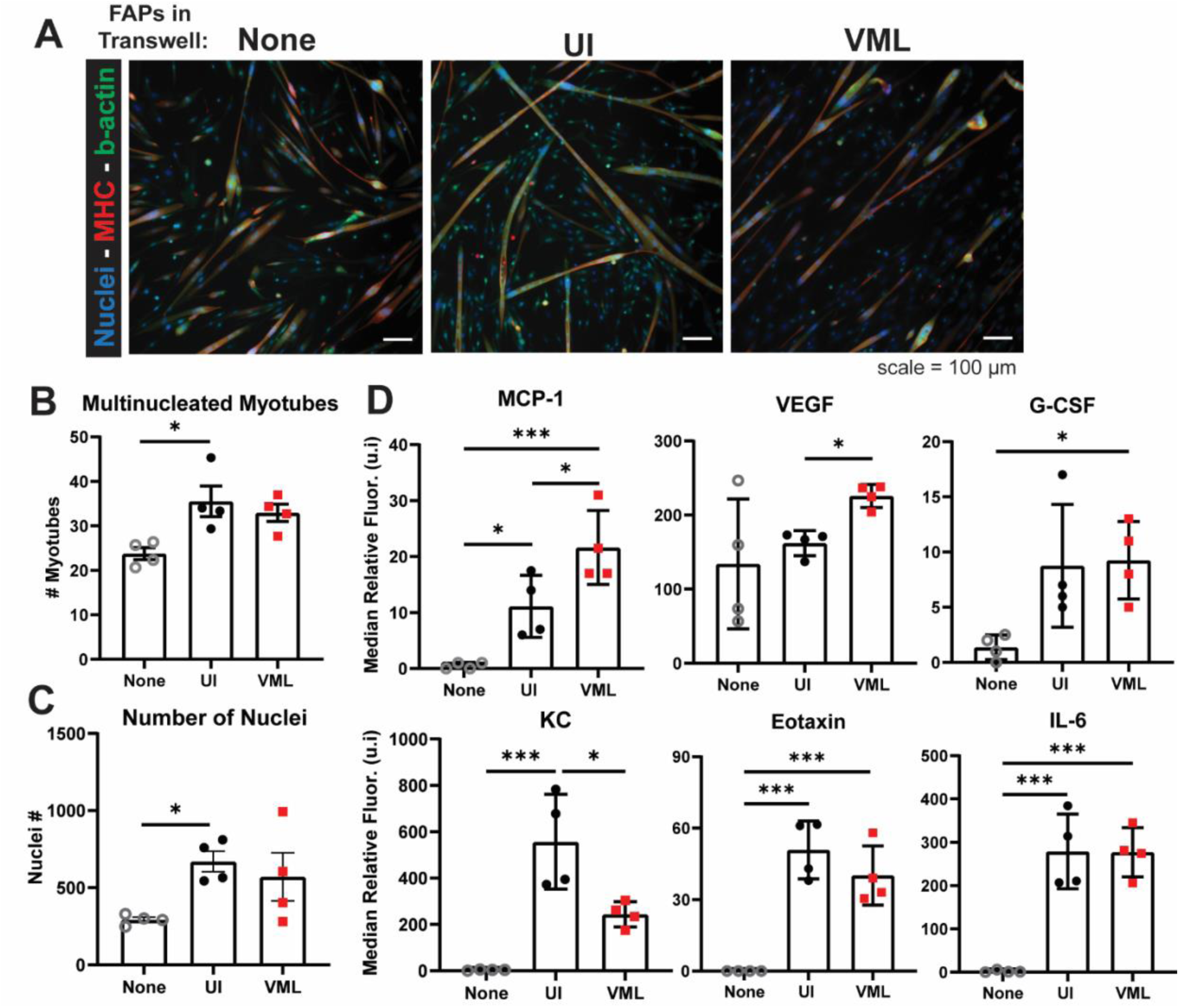
VML derived FAPs do not increase myogenesis and have altered cytokine expression when cultured in transwell with MuSCs. Representative IHC images of GFP-MuSCs cultured alone or following co-culture with uninjured or VML derived FAPs isolated 7 days post-injury in transwell (**A**). MuSCs stained for nuclei (blue), myosin heavy chain (MHC, red), and anti-GFP for amplification of b-actin-GFP signal (green). Multinucleated myotubes (**B**) and total nuclei (**C**) were counted and compared between culture conditions after 8 days of culture. Cytokines present in media at day 8 were measured and detected cytokines are shown (**D**). One-way ANOVA with Tukey’s test post-hoc, *=*p*<0.05, ***=*p*<0.001. n=4 replicates per experimental group.

To measure differences in secreted factors of co-cultured MuSCs and FAPs, we collected culture media from the transwells of each group before fixation at day 8 and measured the cytokine concentrations in each group (Fig. 4D). IL-6 and Eotaxin were significantly elevated with either uninjured or VML derived FAPs in transwell with MuSCs compared to untreated MuSCs in culture alone. Monocyte chemoattractant protein-1 (MCP-1) and vascular endothelial growth factor (VEGF) were significantly increased in the co-culture condition with VML derived FAPs in transwell compared to uninjured FAPs. Additionally, MCP-1 and G-CSF were significantly increased with VML derived FAPs in transwell from the MuSCs cultured alone. KC secreted in the uninjured FAPs transwell condition was significantly elevated above MuSCs alone and co-culture with VML derived FAPs. These differences in cell crosstalk between MuSCs and uninjured versus VML derived FAPs may be important in the progression of fibrosis rather than myogenesis in the injury environment following critical VML.

### Identification of a unique, pro-fibrotic FAPs subpopulation in VML

FAPs post-VML showed increased responsiveness to the pro-fibrotic stimuli TGF-β1 implicating differences in the subpopulation of FAPs present in the VML injury compared to uninjured or regenerating skeletal muscle. Thus, we hypothesized that a subpopulation of FAPs might be contributing to the significant fibrosis following VML. To evaluate this, we performed single cell RNA sequencing (scRNA-seq) to elucidate gene signatures between pro-reparative and pro-fibrotic FAP subpopulations. Cells were isolated from injured quadriceps 7 days post subcritical or critical VML injury as well as from uninjured quadriceps. Single cells from each experimental condition were clustered in an unsupervised approach based on their gene expression profiles and embedded within a low-dimensional uniform manifold approximation and projection (UMAP) for visualization. UMAP plots of the clustering results revealed five cell types comprised in the extracellular matrix (ECM) meta-cluster: tenocytes, FAPs, and three heterogeneous fibroblast subsets (Fig. 5A). Differential gene expression between these five clusters (Fig. S2, A and B) allowed for the identification of these cell types present in the quadriceps across experimental conditions. The identified FAP cluster displayed high expression of *Ly6a*, *Cd34*, and *Pdgfra* (Fig. S2C) which is consistent with previously published literature to validate FAP characterization (*19, 20*).

**Fig. 5.**
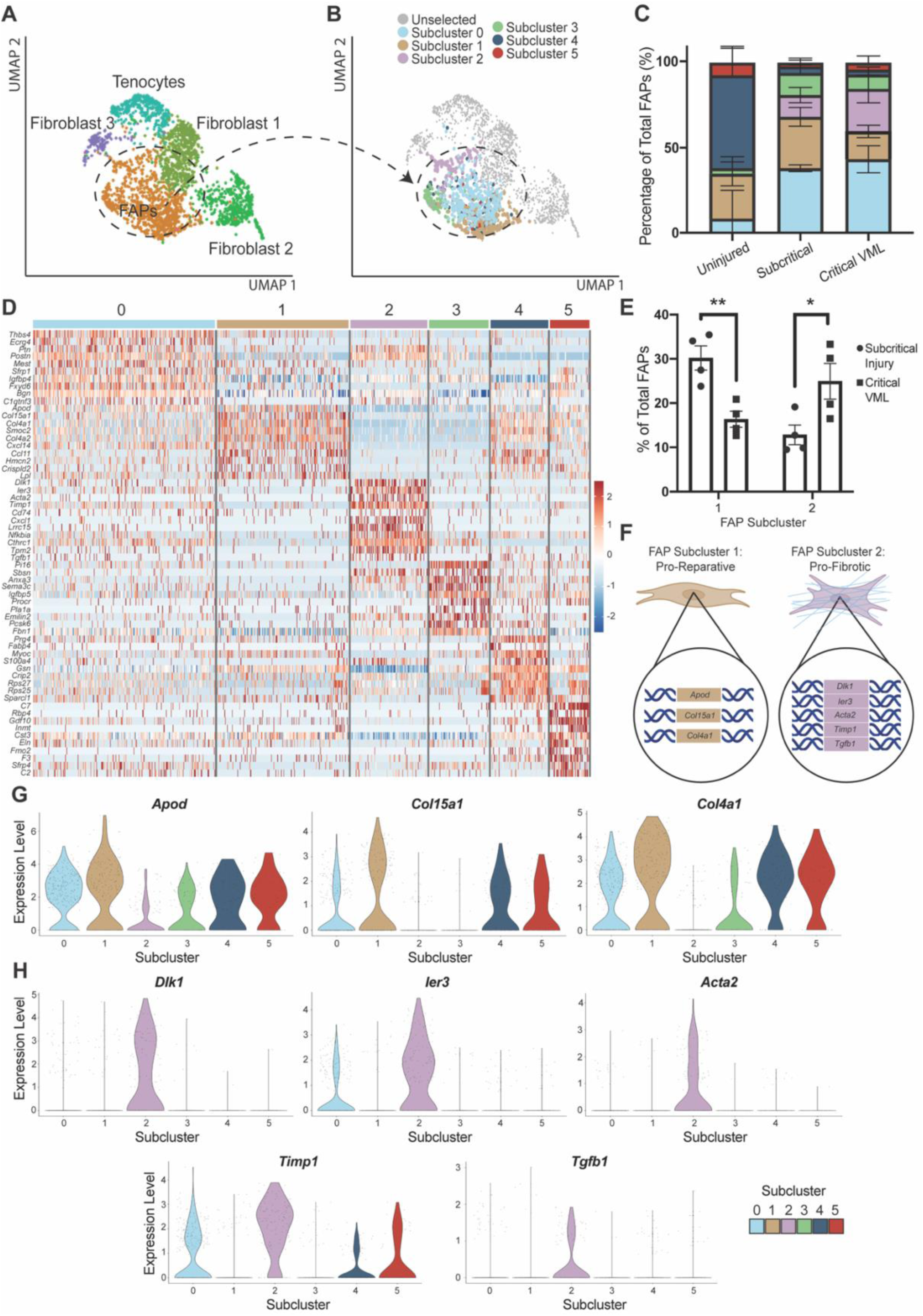
Skeletal muscle from critical VML harbors pro-fibrotic FAP subpopulation with unique RNA signature. (**A**) UMAP representation of identified extracellular matrix cell populations from uninjured, subcritical injury, and critical VML quadriceps at 7 days post injury. (**B**) UMAP overlay of the six identified FAP subclusters determined by differential RNA expression of all FAPs. Tenocyte and fibroblast populations are colored in gray. (**C**) Stacked bar graph quantification of FAPs per subcluster as a percentage of all FAPs for each injury condition. (**D**) Heatmap displaying the top genes expressed per FAP subcluster. Average gene expression denoted by color scale of blue to red, representing low to high expression, respectively. (**E**) Statistical comparison of the number of FAPs belonging to subcluster 1 and 2 between subcritical injury and critical VML injury. (**F**) Schematic representation of the tops genes differentiating pro-reparative FAP subcluster 1 from pro-fibrotic FAP subcluster 2. (**G**) Violin expression plots per FAP subcluster for top genes of FAP subcluster 1: *Apod*, *Col15a1*, and *Col4a1*. (**H**) Violin expression plots per FAP subcluster for top genes of FAP subcluster 2: *Dlk1*, *Ier3*, *Acta2*, *Timp1*, and *Tgfb1*. Data presented as mean ± S.E.M. Statistical analyses performed include two-way ANOVA with repeated measures design. *=*p*<0.05, **=*p*<0.01. n=4 animals per experimental group.

To further explore the discrete heterogeneities underlying FAPs’ differing roles in muscle injury pathology, we sub-clustered the single cells comprising the FAP cluster via unsupervised modularity optimization. Notably, six distinct FAP subsets resulted as determined by their unique transcriptional factors (Fig. 5, B and D). While FAP subcluster 4 and 5 were largely present in uninjured muscle, the transcriptional profile of subsets 0-3 were primarily induced upon muscle injury. FAPs from subcluster 0 made up around 40% of all FAPs in both subcritically injured and critical VML injured muscle (Fig. 5C). This subcluster was characterized by its high expression of thrombospondin 4 (*Thbs4,* Fig. 5D), an ECM protein known to regulate tissue growth and remodeling post-injury (*21*). Interestingly, FAPs from subcluster 1 were significantly increased in subcritically injured muscles compared to critically injured quadriceps (Fig. 5E). This subset presented with elevated levels of Apolipoprotein D (*ApoD*) along with *Col15a1* and *Col4a1* (Fig. 5, D, F and G), both integral collagen components of the basement membrane. In contrast, critical VML resulted in elevated numbers of FAPs from subcluster 2 (Fig. 5E). FAP subcluster 2 is defined by high *Dlk1* expression (Fig. 5, D and H), and overexpression of *Dlk1* is known to inhibit adipogenesis (*22, 23*). Furthermore, this FAP subset highly expresses *Acta2* (Fig. 5H), the gene encoding αSMA, which we showed is increased in VML derived FAPs in the presence of TGF-β1 (Fig. 3D). In addition to *Dlk1* and *Acta2*, FAP subcluster 2 is characterized by its high expression of genes *Tgfb1*, tissue inhibitor of metalloproteases (*Timp1*), and immediate early response 3 (*Ier3*) (Fig. 5, D, F and H).

FAP heterogeneity between injury environments and uninjured skeletal muscle was further characterized by applying UMAP and SPADE analyses to multiparameter flow cytometry data to assess surface marker signatures. UMAP was used to create a low-dimensional space which clustered all FAPs events from uninjured, subcritically, and critically injured muscle tissue at 1, 3, and 7 days post-injury. Events from each group were then overlaid on the UMAP to visualize each groups’ clustering and movement over time. FAPs from uninjured muscle clustered in the bottom left corner of the UMAP (Fig. 6A). Over time after injury overlaid FAPs shift more predominantly towards the upper right quadrant of the UMAP space (Fig. 6B). At 1 and 3 days post-injury, FAPs from subcritical and critical injuries are located in similar locations on the UMAP. However, at 7 days post-injury, subcritical injury FAPs shift back to the left of the UMAP, but critical VML FAPs remain on the right. Critical VML FAPs are also present at qualitatively higher densities on the UMAP than subcritical injury derived FAPs, illustrating the previously quantified difference in concentration (Fig. 1). The surface marker with the strongest correlation to the observed temporal UMAP shift of VML derived FAPs was β1-integrin. β1-integrin expression overlaid onto the UMAP projection shows that its expression increases in the same pattern as movement of critical VML derived FAPs over time, from bottom left to top right of the UMAP (Fig. 6C).

**Fig. 6.**
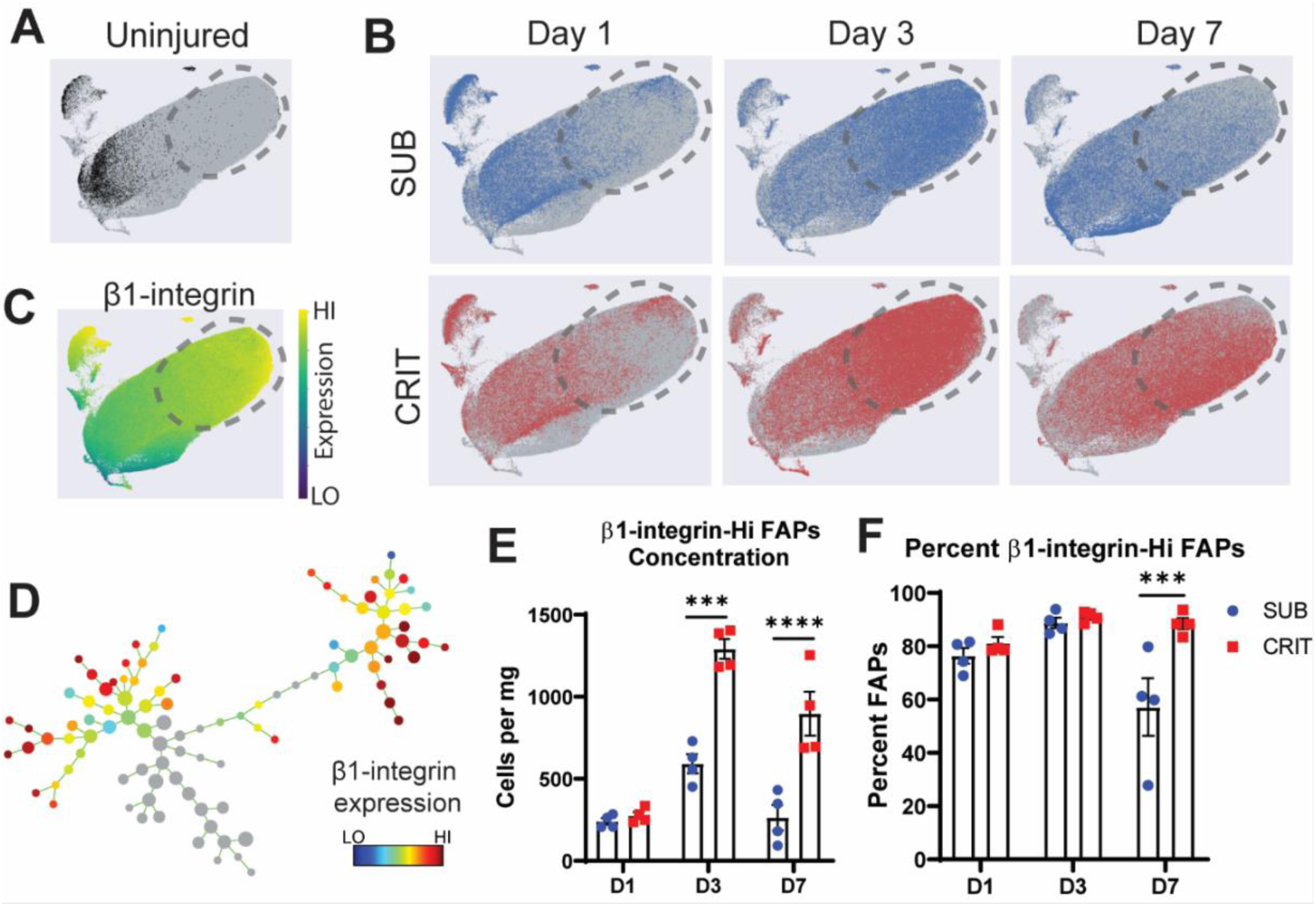
Dimensionality reduction techniques UMAP and SPADE identify a subpopulation of FAPs in critically injured muscle which highly express β1-integrin. A UMAP was generated for all FAPs analyzed via flow cytometry and overlaid with events from each group, uninjured muscle (**A**, UI, black), subcritical injured (SUB, blue) and critical injured (CRIT, red) (**B**). Grey dotted line drawn on after to illustrate the β1-integrin high region, shown by the surface marker expression overall (**C**). A SPADE tree was generated (**D**) from the same flow cytometry data used for the UMAP. Colored nodes were considered β1-integrin highly expressing and quantified as concentration (cells/mg, **E**) and percentage total FAPs (**F**). Statistics performed in GraphPad Prism 8, two-way ANOVA with Sidak test post-hoc, ***=*p*<0.001, ****=*p*<0.0001. n=4 animals per experimental group.

A SPADE tree was generated to quantify the concentration and relative percentage of β1-integrin high expressing (β1-HI) FAPs in skeletal muscle from subcritical injury and critical VML injuries (Fig. 6D). Nodes shown in color were considered β1-HI and used for quantification. The number of FAPs per tissue mass was quantified, showing a significantly greater concentration of FAPs in critically injured tissue at 3 and 7 days post-injury were β1-HI compared to subcritically injured muscle (Fig. 6E, *p*<0.001). The percentage of total FAPs which were β1-HI was quantified; the proportion of β1-HI FAPs in critical VML tissue remained elevated at a level which was significantly higher than in subcritical injury tissue (Fig. 6F, *p*<0.001). As persistent fibrosis is a hallmark of critical VML, this population of FAPs which was specifically increased following critical injury warranted further investigation as a potential pro-fibrotic subpopulation.

FAPs at 7 days post-injury from critically injured VML tissue were sorted based on their β1-integrin expression to assess the role of β1-integrin in driving fibrosis. Isolated β1-HI and β1-integrin low expressing (β1-LO) cells were then used to determine differences in gene expression and *in vitro* differentiation. A mouse fibrosis quantitative PCR (qPCR) array was used to quantify differences in pro-fibrotic gene expression in uninjured, β1-HI, or β1-LO FAPs. Key genes from this fibrotic qPCR array were upregulated in critical VML-enriched FAP subcluster 2 from our scRNA-seq subpopulation analysis relative to pro-reparative subcluster 1 (Fig. 7A), including significant elevation of *Itgb1* (the gene for β1-integrin). These data further suggest that the unsupervised identification of FAP subcluster 2 in VML is implicit in the pathological fibrotic response. These key fibrotic genes were also crucial in differentiating FACS isolated β1-HI from β1-LO FAPs from critically injured tissue (Fig. 7B) which indicates that β1-HI FAPs are likely derived from this transcriptionally distinct subcluster. PLSDA, a supervised clustering technique, was used to look for differences between the isolated FAPs in a two-dimensional projection. Uninjured, β1-HI, and β1-LO FAPs separated along two LVs, with injured versus uninjured separating along LV1 and β1-HI versus β1-LO separating along LV2 (Fig. 7C). The relative contributions of each gene to each latent variable can give indication of the meaning underlying separation along LV1 and LV2 in the PLSDA plot (Fig. 7, D and E). β1-HI and β1-LO FAPs had significantly higher LV1 scores than uninjured FAPs (Fig. 7F, *p*<0.01). On LV2, β1-HI FAPs had significantly higher scores than both uninjured and β1-LO FAPs (Fig. 7G, *p*<0.05). Genes for collagen 3 (*Col3a1*), *Timp1*, thrombospondin 1 (*Thbs1*), and β1-integrin (*Itgb1*) contributed to higher scores on LV1 and LV2; suggesting these genes may be upregulated post-injury and with β1-integrin expression. On LV1 specifically, the *Tnf* gene was associated with low scores, indicating that uninjured FAPs express more TNF than VML derived FAPs. On LV2 specifically, the genes for αSMA (*Acta2*) and TGF-β1 (*Tgfb1*) were both associated with higher scores and therefore with β1-HI FAPs. Conversely, *Mmp3* was one gene associated with β1-LO FAPs specifically. Similarly, separation among FAPs groups were observed with unsupervised principal component analysis (PCA) with uninjured, β1-HI, and β1-LO FAPs separating along principal component 1 (PC1, Fig. S3A). Scores for each sample on PC1 and PC2 were compared (Fig S3, B and C) and scores for PC1 significantly increased from uninjured to β1-LO to β1-HI. The corresponding loading plot for PC1 (Fig. S3D) revealed similar genes of interest for β1-HI FAPs to those from PLSDA including *Timp1*, *Acta2*, and *Tgfb1*.

**Fig. 7.**
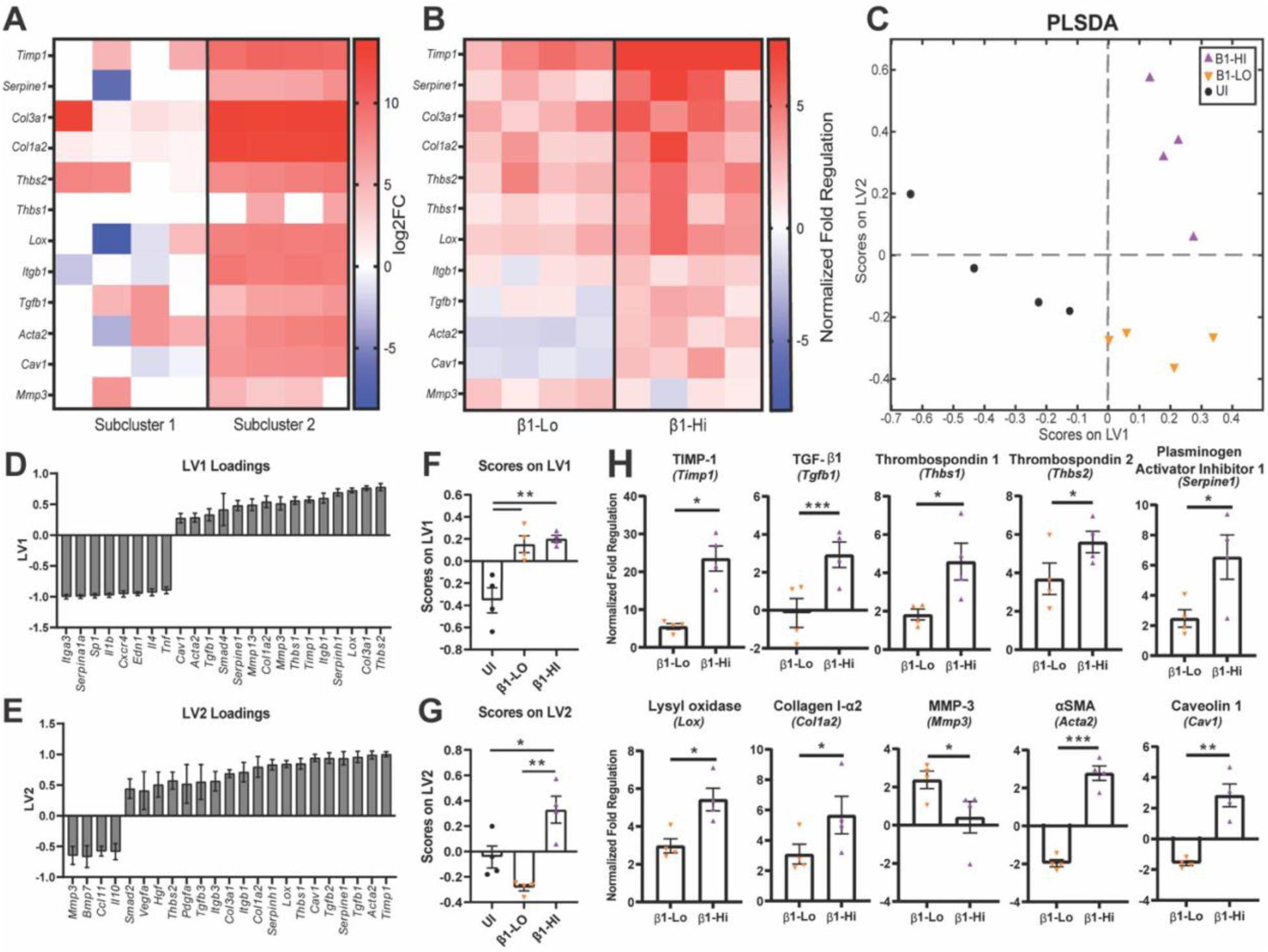
β1-HI FAPs present with a distinguished pro-fibrotic gene signature. Single-cell RNA expression of selected fibrotic genes for pro-reparative FAP subcluster 1 and pro-fibrotic subcluster 2 presented as a heatmap (**A**). Expression quantified as log2 fold change between critical VML (CRIT) and uninjured (UI). Heatmap showing relative gene expression from qPCR for FAPs isolated from UI or CRIT tissue segmented into β1-integrin high expressing (β1-HI) and β1-integrin low expressing (β1-LO) (**B**). Data quantified as normalized fold regulation. PLSDA plot of UI, β1-HI, and β1-LO FAPs according to their scores on two latent variables (LV) 1 and 2 (**C**). Select high and low gene loadings on LV1 and LV2 (**D** and **E**) and LV scores between conditions (**F** and **G**) were plotted and compared. Significantly different genes are plotted (**H**) and labelled as “Protein Name (Gene Name)”. One-way ANOVA with Tukey’s test post-hoc, paired student’s t-test performed for biologically matched samples, *=*p*<0.05, **=*p*<0.01, ***=*p*<0.001, ****=*p*<0.0001. n=4 animals per experimental group.

As clustering methods were able to separate uninjured, β1-HI, and β1-LO FAPs, we then directly compared the expression levels of specific genes. Values are reported as normalized fold regulation relative to uninjured FAPs. Gene expression levels of each sample were visualized in heatmaps (select genes, Fig. 7B, all genes, Fig. S3E) and those genes with significant differences were compared (Fig. 7H, *p*<0.05). *Tgfb1*, the gene for the pro-fibrotic growth factor TGF-β1, was significantly elevated in β1-HI versus β1-LO FAPs (*p*<0.001). *Thbs1* encodes thrombospondin 1 which can activate latent TGF-β1 and is also upregulated in β1-HI compared to β1-LO FAPs (*p*<0.05). *Timp1*, *Col1a2*, *Serpine1*, *Thbs2*, and *Lox* were all significantly upregulated in β1-HI compared to β1-LO FAPs (*p*<0.05). These genes are for proteins which collectively contribute to ECM deposition and cross-linking. *Acta2* and *Cav1* were also significantly elevated in β1-HI FAPs compared to β1-LO FAPs (*p*<0.01) and are related to cell structure. *Acta2*, specifically, encodes αSMA, a marker of myofibroblast differentiation (used in Fig. 3 IHC). The only gene which was significantly downregulated in β1-HI FAPs was a gene for ECM degradation, *Mmp3*. The parallels in gene signatures indicate that our scRNA-seq identified pro-fibrotic FAPs (subcluster 2) likely yield β1-HI FAPs in critically injured tissue at the surface marker level. These results suggest that β1-HI FAPs may functionally promote a pro-fibrotic environment.

### TIMP1 and TGF-β1 are implicated in the promotion of fibrosis and inhibition of myogenesis in critical VML

To further investigate the β1-HI subpopulation of FAPs following VML, we cultured β1-HI and β1-LO from PDGFRα^EGFP^ mice *in vitro*. The goal of these experiments was to determine whether VML derived FAPs would have altered cytokine secretion or differentiation potential dependent on β1-integrin expression. The concentration of TGF-β1 in the culture media was quantified after 6 days in growth media; there were no significant differences in TGF-β1 levels in the media between β1-HI and β1-LO FAPs (Fig. 8A). However, when stimulated with TGF-β1, β1-HI FAPs expressed more αSMA than the unstimulated control, indicating increased myofibroblast differentiation (Fig. 8, B and C, *p*<0.01). Conversely, β1-LO FAPs did not show a significant increase in αSMA when treated with TGF-β1 (Fig. 8B, *p*>0.05).

**Fig. 8.**
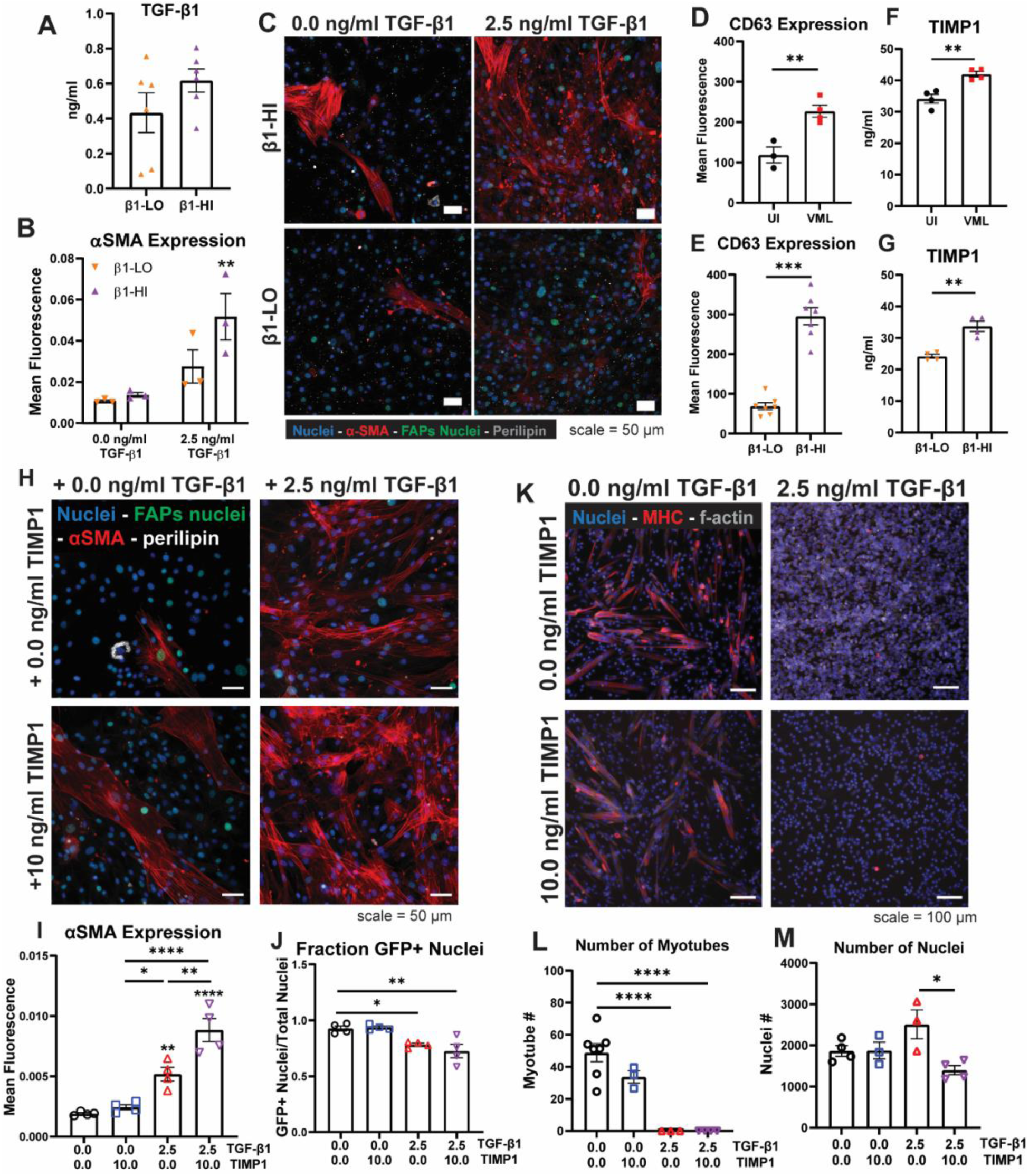
TGF-β1 and TIMP1 synergistically inhibit myogenesis. The concentration of TGF-β1 present in media at day 6 in culture was measured (**A**) immediately before the differentiation was started. Representative images after differentiation with or without TGF-β1 stimulus (**C**) IHC staining performed for αSMA (red), nuclei (blue), FAPs nuclei (green), and perilipin (grey). αSMA fluorescence was quantified (**B**). Mean fluorescence intensity of CD63 staining as measured by flow cytometry of uninjured (UI) and critical VML (VML) FAPs (**D**) or of β1-HI and β1-LO FAPs (**E**). TIMP1 concentration in conditioned media from UI and VML (**F**) or β1-HI and β1-LO FAPs (**G**). Representative IHC images of UI FAPs cultured with either TGF-β1, TIMP1, neither, or both (**H**) with staining for nuclei (blue), FAPs nuclei (green), αSMA (red), perilipin (grey). Mean αSMA fluorescence (**I**) and the fraction of GFP^+^ nuclei (**J**) were quantified. Representative images of MuSCs cultured with TGF-β1, TIMP1, neither, or both following fixation and IHC staining (**K**) stained for nuclei (blue), myosin heavy chain (MHC, red), and f-actin (grey). The number of multinucleated myotubes (**L**) and nuclei (**M**) were quantified. Two-way ANOVA, Sidak test post-hoc (**B**), t-tests (**D** to **G**), one-way ANOVA with Tukey’s test (**I** and **M**) or Dunnett’s test (**J** and **L**) post-hoc, *=*p*<0.05, **=*p*<0.01, ***=*p*<0.001, ****=*p*<0.0001. n=3-7 replicates per experimental group.

From the qPCR and *in vitro* differentiation analyses, β1-HI FAPs from the VML injuries were a highly pro-fibrotic subset of FAPs. While TGF-β1 has been shown here and in previous studies to be a clear pro-fibrotic stimulus in FAPs culture (*6*), the stark increase in β1-HI TIMP1 gene expression (Fig. 7H) drove us to investigate whether TIMP1 played a role in FAPs differentiation. While TIMP1 is typically studied for its role in inhibiting the degradation of ECM proteins through the enzymatic action of matrix metalloproteinases (MMPs), it has recently been studied for its ability to function as a signaling molecule (*24*). In order to impact intracellular cell signaling, TIMP1 binds a surface receptor complex of β1-integrin and CD63 (*25*).

Surface expression of CD63 was measured via flow cytometry on the surface of FAPs which were from either uninjured or critical VML injured tissue. CD63 expression was significantly upregulated in VML versus uninjured FAPs (Fig. 8D, *p*<0.01). Additionally, FAPs from uninjured and VML tissue gated on β1-integrin expression showed a significant increase in CD63 fluorescence in β1-HI FAPs (Fig. 8E, *p*<0.001). TIMP1 levels were measured in the cell culture media of uninjured or VML derived FAPs at day 6 in culture; VML derived FAPs were found to secrete significantly increased levels of TIMP1 (Fig. 8F, *p*<0.01). VML-derived FAPs which were sorted based on β1-integrin expression also had significantly different TIMP1 secretion *in vitro*, with β1-HI FAPs secreting more TIMP1 than their β1-LO counterparts (Fig. 8G, *p*<0.001).

To compare the pro-fibrotic signaling capabilities of TIMP1 to TGF-β1, uninjured FAPs were cultured for 6 days before beginning differentiation in media containing TGF-β1, TIMP1, or both. IHC staining was done to measure FAPs differentiation (Fig. 8H) and αSMA expression increased significantly when FAPs were stimulated with TIMP1 and TGF-β1 over TGF-β1 alone (Fig. 8I, *p*<0.01). TIMP1 alone did not significantly increase αSMA expression (Fig. 8I). TGF-β1 with or without TIMP1 reduced the fraction of PDGFRα-GFP^+^ nuclei similarly when compared to unstimulated FAP controls (Fig. 8J).

While it has been characterized that β1-integrin is required for TIMP1 signaling, β1-integrin has also been shown to play a role in kidney and liver fibrosis in the context of TGF-β1 stimulation (*26, 27*). We therefore wanted to investigate whether inhibiting binding to β1-integrin using a blocking antibody (anti-ITGB1) would decrease the fibrotic differentiation of FAPs stimulated with TGF-β1. Following FAPs differentiation with TGF-β1 with or without anti-ITGB1 supplemented in the media, the mean αSMA fluorescence was quantified (Fig. S4, A to C). FAPs cultured with TGF-β1 had a significant increase in αSMA fluorescence, consistent with previous experiments (Fig. S4D, *p*<0.05). However, when treated with both TGF-β1 and anti-ITGB1, FAPs did not have a significant increase in fibrotic differentiation compared to the untreated control (Fig. S4D, *p*>0.05). These results, consistent with findings in tissue fibrosis of other organs, provide a potential target for reducing fibrotic differentiation of FAPs following critical VML.

Following characterization of TIMP1 and TGF-β1 as pro-fibrotic signaling molecules secreted by β1-HI FAPs, we investigated whether these cytokines may affect myoblast differentiation. MuSCs were cultured with TGF-β1, TIMP1, neither, or both (Fig. 8K), as was done with FAPs (Fig. 8H). MuSCs cultured with TIMP1 alone did not cause a significant change in the number of multinucleated myotubes compared to untreated MuSCs (Fig. 8L, *p*>0.05). However, MuSCs cultured with TGF-β1 were unable to fuse to form any multinucleated myotubes, a significant reduction compared to the untreated control (Fig. 8L, *p*<0.0001). When comparing the number of nuclei, neither TGF-β1 nor TIMP1 alone caused a significant change in nuclei from the untreated control. However, when TGF-β1 and TIMP1 were added concurrently, there was a significant reduction in the number of nuclei compared to the TGF-β1 alone treated MuSCs (Fig. 8M, *p*<0.05). Taken together, these data may indicate that TGF-β1 and TIMP1 could be working to slow MuSC proliferation and differentiation following VML.

## Discussion

VML results in substantial fibrosis rather than functional muscle regeneration. While FAPs have been identified as a source of fibrosis in muscle myopathies, there is a poor understanding of their role in the context of VML. In these experiments we have laid the groundwork for understanding the interplay between FAPs and MuSCs following a critical VML injury. While there is a high overall concentration of MuSCs and evidence of myoblast fusion events following critical injury, we show that the injury environment has an abundance of pro-fibrotic FAPs that secrete signaling molecules which are simultaneously pro-fibrotic and anti-myogenic. These data implicate a role of β1-integrin and TIMP1 in the chronic fibrotic process and therefore may be novel targets following VML to reduce pro-fibrotic FAPs and enhance muscle regeneration.

In this series of studies, we show that FAPs concentration increased in muscle over time from critical VML injuries and FAPs nuclei made up the majority of nuclei within the defect site. The data indicate that FAPs may play an important role at the threshold of the switch from recoverable (subcritical) to irrecoverable (critical) VML injury. The abundance of FAPs within a site of insufficient muscle regeneration and prolonged immune cell presence led us to ask questions about the role of FAPs’ paracrine signaling following VML, as systemic cytokines may contribute to exacerbated fibrosis and inflammation characteristic of severe injury.

Importantly, we found that KC, VEGF, MCP-1, and TGF-β1 were significantly elevated in FAPs isolated from VML injured muscle compared to uninjured controls. TGF-β1 is considered the main driver of myofibroblast differentiation in FAPs (*28*), and KC has been linked to both neutrophil trafficking (*29, 30*) and the promotion of tissue fibrosis (*31, 32*). These results indicate the VML-derived FAPs are biased towards a pro-fibrotic and pro-inflammatory phenotype relative to their quiescent phenotype. Following the induction of fibrotic differentiation with or without TGF-β1 supplementation, we found that IL-6 and MCP-1 were significantly elevated in VML FAPs treated with TGF-β1 while cytokine levels in uninjured FAPs remained constant regardless of TGF-β1 supplementation. These results highlight that VML FAPs present with increased responsiveness to TGF-β1. In the context of chronic inflammation and fibrosis, MCP-1 and TGF-β1 have been shown to stimulate expression of one another via positive crosstalk, resulting in excessive inflammation and ECM deposition (*33*). FAPs may be an additional source of MCP-1 in VML injured muscle, which may feed forward the recruitment of immune cells and TGF-β1 within the muscle and ultimately impair muscle regeneration.

We found that IP-10, VEGF, and LIF not only significantly increased in VML FAPs once treated with TGF-β1 but was significantly elevated between TGF-β1-treated VML FAPs and TGF-β1-treated FAPs from uninjured muscle. This suggests that TGF-β1 supplementation alone is not responsible for the altered secretory profile of these cytokines, but their tissue-resident niche itself substantially influences their paracrine function. VEGF, a cytokine which is a crucial promoter of angiogenesis, has been shown to stimulate fibrotic differentiation by increasing αSMA and *Col1a1* in liver cells *in vitro* (*34*). Further, VEGF has been shown to play a role in the maintenance of MuSC quiescence (*35*); thus, future studies probing whether VEGF secretion by FAPs elicits an anti-myogenic effect of MuSCs is of interest. Both IP-10 and LIF have been associated with reduced fibrosis and TGF-β1 expression (*36, 37*). While counterintuitive, it is possible that differentiation to myofibroblasts stimulates the release of IP-10 and LIF as a compensatory mechanism which under normal healing circumstances could properly limit tissue fibrosis. However, chronic inflammatory conditions, such as VML, are characterized by an upregulation in both traditionally inflammatory and anti-inflammatory cytokines. Therefore, the precise role of IP-10 and LIF in the context of critical VML will require further investigation.

Most notably, our studies utilize cutting-edge single cell technologies to expand the knowledge that FAPs present with plasticity and heterogeneity dependent on the microenvironment in which they encounter. Traditionally observed as a single progenitor population, only recently have FAPs been highly resolved into distinct subsets dependent on differentiation potential and unique functional role in muscle injury (*38*). Our single-cell RNA sequencing revealed that *Apod*^+^ pro-reparative FAPs localized within recoverable, subcritical injuries. Recent literature shows that FAP-like adipose stromal cells from subcutaneous adipose tissue are crucial to muscle regeneration following injury, thereby suggesting a role for *Apod*^+^ FAPs in regeneration (*39*). In contrast, VML injured muscle presented with significantly increased FAPs co-expressing pro-fibrotic genes *Tgfb1*, *Acta2*, and *Timp1*. Identified pro-fibrotic FAPs from VML injuries highly expressed *Ier3* which functions to protect cells from TNF-mediated apoptosis. As FAP apoptosis is crucial to regulating their proliferation after injury, these results suggest that VML FAPs are primed for fibrotic differentiation rather than regulated apoptosis.

Our multidimensional flow cytometry approach allowed us to identify and sort a distinct FAP subset upregulated in VML with a highly pro-fibrotic gene and protein signature. Leveraging dimensionality reduction techniques, we discovered that β1-integrin was differentially expressed within VML FAPs, and β1-integrin high expressing FAPs (β1-HI) were characterized by increased expression of TIMP1, TGF-β1, *Col1a2*, αSMA, and *Lox* among others compared to β1-LO FAPs. PLSDA and PCA showed separation of FAPs based on β1-integrin expression. These clustering algorithms indicated that *Tgfb1* and *Timp1* gene expression was enriched in the β1-integrin high subpopulation. This common observation of upregulated TIMP1 in addition to TGF-β1 in a specific subset of β1-HI FAPs is one which has not been previously characterized. TIMP1 has been primarily characterized for its activity as an inhibitor of MMPs, which can limit cell migration and exacerbate fibrosis by preventing the degradation and remodeling of ECM proteins (*40*). Recently, TIMP1 has also been found to work synergistically with TGF-β1 in the development of myocardial fibrosis and increased collagen deposition (*41*). To function as a signaling molecule, TIMP1 binds to a CD63 and β1-integrin surface receptor complex (*25*). We found that β1-integrin highly expressing FAPs were present at significantly higher concentrations specifically in critical compared to subcritical injuries. Additionally, we saw increased CD63 surface expression in VML derived FAPs, which was associated with increased β1-integrin expression.

In addition to promoting a fibrotic ECM, β1-HI FAPs also had elevated gene expression for two intracellular proteins. *Acta2a*, the gene for αSMA, and *Cav1*, the gene for caveolin (Cav)-1 were both significantly elevated in β1-HI FAPs over β1-LO FAPs. It has been shown previously that as FAPs differentiate to myofibroblasts, expression of αSMA also increases, as was replicated in the current work. Therefore, high expression of αSMA in undifferentiated β1-HI FAPs reinforces their bias to differentiate down a fibrogenic lineage. Cav-1, a protein which localizes at the cell membrane, has a number of diverse functions, as it is known that Cav-1 plays an important role in focal adhesions and has been found to co-localize with β-integrins (*42*). Through this mechanism, an increase in Cav-1 in β1-HI FAPs may be aiding the increased levels of β1-integrin to form focal adhesions with the surrounding ECM. Together, these data suggest an intracellular environment in which the cytoskeleton and cell membrane of β1-HI FAPs is stabilizing, adhering, and differentiating into myofibroblasts.

β1-integrin has been shown to be a receptor for multiple ligands, not only TIMP1 but also TGF-β1. TGF-β1 is found *in vivo* in a latent form, bound to latency associated proteins (LAPs). These LAPs have RGD-peptide sequences and have been shown to bind to various integrins, including β1-integrin with αV and α8, which could serve to localize TGF-β1 near the surface of β1-HI FAPs (*43, 44*). Our results indicate that blocking β1-integrin reduced TGF-β1 induced myofibroblast differentiation of FAPs *in vitro*, however the mechanism of this phenomenon was not determined. Whether there was an effect blocking TIMP1 binding, directly interfering with TGF-β1 signaling, or impairment of TGF-β1 activation are both potential explanations. We not only found that TGF-β1 and TIMP1 synergistically enhance FAP fibrotic differentiation, but additionally, the combination negatively impacted MuSC proliferation. These findings suggest that β1-integrin and its downstream signaling pathway could be a target for future therapeutic strategies for VML-related fibrosis. While our studies elucidate the heterogeneity of FAPs driving fibrosis, the role of FAPs in fatty infiltration following VML remains underexplored. A recent study found that FAPs are the primary source for fatty infiltration in full-thickness rotator cuff tear in humans but are capable of adopting a pro-myogenic beige adipose tissue phenotype (*45*). Thus, investigating the plasticity of FAPs in VML-induced fatty infiltration will be an intriguing focus of future studies.

In summary, our results are foundational in understanding the dynamic response of MuSCs and FAPs and their effect on regeneration following critical VML injury in the mouse quadriceps. Our work highlights that uninjured or subcritical injuries present with a FAP subset population that distinctly differs from VML-derived FAPs at the transcriptional, protein, and secretome level. This work implicates a novel, aberrant pro-fibrotic FAP subpopulation as a driver of persistent fibrosis and inflammation which impairs myogenesis in VML. These studies harnessing in depth single-cell profiling of heterogeneous muscle progenitor populations can be utilized to inform and direct future therapeutic targets for VML-related fibrosis.

## Materials and Methods

### Animals

B6.129S4-*Pdgfra^tm11(EGFP)Sor^*/J (PDGFRα^EGFP^), in which cells containing the surface marker PDGFRα have enhanced green fluorescent protein (EGFP) expressing nuclei, were purchased from Jackson Laboratory and used for histological quantification and cell sorting and *in vitro* culture experiments. Male and female adult mice 4-7 months old were used. Pax7^+^ MuSC reporter (Pax7TdT) mice were bred by crossing *B6.Cg-Pax7tm1(cre/ERT2)Gaka/J* and B6.Cg-*Gt(ROSA)26Sor^tm9(CAG-tdTomato)Hze^*/J mice from Jackson Laboratories. The resulting Pax7TdT mice expressed TdTomato red fluorescent protein in Pax7^+^ cells following tamoxifen treatment. C57BL/6-Tg(CAG-EGFP)1Osb/J (b-actin GFP) mice expressing green fluorescent protein (GFP) under control of the b-actin promoter were purchased and bred for utilization in MuSC-FAPs transwell culture experiments. C57BL/6J mice were purchased from Jackson Laboratories and were used for cell sorting, *in vitro* culture experiments, and subcritical versus critical injury characterization. Adult male mice, referred to as “young”, were on average 6.1±0.5 (mean ± standard deviation) months in age at the time of euthanasia. All animals were used according to the protocols approved by the Georgia Institute of Technology’s IACUC.

### Volumetric muscle loss injury

Surgical procedure performed as previously reported (*15*). Briefly, mice were anesthetized with 2% isoflurane, the left hindlimb was prepped and sterilized. A single incision was made above the quadriceps and a 2mm (subcritical) or 3mm (critical VML) biopsy punch (VWR, 21909-132, −136) was used to make a full-thickness muscle defect. Skin was closed and muscle was left to recover without intervention for 1, 3, 7, or 14 days before euthanasia by CO_2_ inhalation.

### Tissue histology and immunostaining

Tissue processing and histology done as previously reported (*15*). Briefly, muscle was dissected, weighed, and snap frozen in liquid nitrogen cooled isopentane. 10 µm cryosections were taken. Samples were blocked and permeabilized before staining with Alexa Fluor 647 phalloidin (ThermoFisher, A22287, 1:250) and Alex Fluor 488 conjugated anti-GFP (ThermoFisher, A21311, 1:250) for 1-hour incubation at room temperature. Slides were mounted with Fluoroshield Mounting Medium with DAPI (Abcam, ab104139) and stored at 4°C.

### Whole mount imaging

Hindlimbs were fixed with muscles attached in 4% PFA for 45 minutes, washed with PBS, and kept in PBS at 4°C until dissection. Quadriceps were dissected and incubated in 20% w/v sucrose overnight at 4°C. Quadriceps were washed in PBS, dried, and pinned to a polydimethylsiloxane (PDMS) filled petri dish. Two razor blades were placed on the lateral and medial sides of the muscle and a scalpel was used to create longitudinal sections. Sections were stained in microcentrifuge tubes on a rotating rack.

### Imaging and quantification of percent GFP^+^ nuclei

Images were taken at 20x magnification using a Zeiss fluorescent microscope. Images were imported into ImageJ/FIJI, converted to 8-bit, thresholded using “Moments” built in method, and then analyze particles. This was done for GFP and DAPI channels. The area occupied by particles was summed for “GFP occupied area” metric and number of GFP^+^ particles was then divided by number of DAPI^+^ particles to give “Percent GFP^+^ nuclei” metric.

### Enzyme-linked immunosorbent assays (ELISAs) and Luminex multiplex cytokine assays

Media was collected at either 6 or 10 days in culture, centrifuged at 10,000xg for 5 minutes. Supernatant was collected and frozen at −80°C. MILLIPLEX MAP Mouse Cytokine/Chemokine Magnetic Bead Panel - Immunology Multiplex Assay (32-plex mouse cytokine assay, Millipore Sigma, MCYTOMAG-70K) was used with for cytokine/chemokine multiplex measurements according to kit instructions. TGF-β1 (ThermoFisher, BMS608-4) and TIMP1 (ThermoFisher, EMTIMP1) ELISAs were used according to manufacturer instructions.

### Single cell RNA sequencing of injured quadriceps

Subcritical injury and critical VML defects were created as described, and animals were euthanized 7 days post injury. Uninjured quadriceps were used as naïve controls. Single cells from quadriceps were isolated as described previously (*46*). Red blood cells were lysed (RBC lysis buffer, BioLegend) for 15 minutes at room temperature, samples were centrifuged, and pelleted cells were washed with cell staining buffer (BioLegend) after supernatant was aspirated. Cells were counted so that 125,000 cells per sample were transferred to new tubes, centrifuged, and resuspended with Fc blocking antibody in cell staining buffer for 10 minutes. Cells from each sample were subsequently stained with TotalSeq-C cell hashtag reagents (BioLegend) to barcode cells from individual animals. Samples from animals in the same experimental group (n=4) were then pooled to total 500,000 cells per condition (3 conditions: uninjured, subcritical, critical VML). Cells were washed and resuspended to a final concentration of 1,000 cells/μL. From here, the 10x Genomics Chromium Next GEM Single Cell 5’ Reagent Kits v2 (Dual Index) protocol was followed to complete single cell preparation for sequencing (CG000331). In brief, ∼10,000 cells (per 500,000 pooled condition group) were loaded onto Chromium Next GEM Chip K for Gel Beads-in-emulsion (GEM) generation. After GEM generation, Gel Beads were dissolved, and co-partitioned single cells were lysed using a thermal cycler to produce 10x Barcoded, full-length cDNA from mRNA. GEMs were broken and pooled following cDNA synthesis, and 10x Barcoded cDNA was purified and subsequently amplified according to protocol. Quality control and quantification of cDNA was analyzed on an Agilent Bioanalyzer High Sensitivity chip and Qubit fluorometer. 5’ gene expression libraries were constructed from amplified cDNA, and sample indices were added to allow for simultaneous sequencing of multiple samples in accordance with 10x Genomics protocol. Post library construction quality control was validated with Agilent Bioanalyzer High Sensitivity chip and Qubit fluorometer. Samples were sequenced on an Illumina sequencer using the NextSeq Mid Output (150 cycles) kit. BCL files were demultiplexed to FASTQ files using Illumina’s “bcl2fastq” function. FASTQ files were aligned to the GRCm39/mm39 mouse reference genome and converted to count matrices using Cell Ranger’s “count” function. Count matrices were then read into and processed with Seurat in R studio.

### Flow cytometry and fluorescence activated cell sorting (FACS)

Samples were prepared for flow cytometry analysis on a FACS AriaIII flow cytometer (BD Biosciences) as previously reported (*46*). Entire injured, left quadriceps were harvested and digested with 5,500U/ml collagenase II and 2.5U/ml Dispase II for 1.5 hours on a shaker at 37°C. Muscle digest solution was filtered through a cell strainer. Single-cell suspensions for flow cytometry were stained for live cells using Zombie NIR (Biolegend) dyes in cell-culture grade PBS per manufacturer instructions before fixation with 4% paraformaldehyde (PFA). If cells were being sorted, antibody staining immediately followed filtration. Cells were stained with cell phenotyping antibodies. The following antibodies we used: FITC-conjugated anti-Ly6A/E (BioLegend), APC-conjugated Lineage antibody cocktail (BD Pharmigen), APC-conjugated anti-CD31 (BioLegend), PE-Cy5 conjugated anti-CD29 (BioLegend), and PerCP-Cy5.5-conjugated anti-CXCR4 (BioLegend). 30mL of CountBright Absolute Counting Beads (C36950, Invitrogen) were added per sample for absolute quantification of cell populations. For FACS, propidium iodide (PI) was added immediately before sorting for viability. Cells were sorted into whole cell growth media supplemented with b-FGF at 2.5 ng/ml. If FAPs sorting on β1-integrin expression, “β1-integrin-Hi” was considered to be the 30% highest expressing FAPs and “β1-integrin-Lo” were considered the lowest 30% expressing FAPs.

### Fibrosis PCR Array

FAPs were isolated by FACS as described above from uninjured (UI) animals as controls, experimental groups were β1-integrin-Hi and β1-integrin-Lo expressing VML-derived FAPs. Immediately following sorting, cells were centrifuged at 300xg for 5 minutes at 4°C, supernatant was removed. Cells were lysed with 1% 2-Mercapto-ehtanol in RNeasy Plus lysis buffer (Qiagen, 1030963) and freeze-thawed 3x in liquid nitrogen. Final thawed solution was centrifuged at 18,000xg for 10 minutes at 4°C. Supernatant was collected and stored at −80°C until RNA purification. RNA purification was completed using RNeasy Mini Kit (Qiagen, 74104) according to kit instructions. RNA was diluted in RNase-Free Water (Qiagen, 1017979) normalized to cell number collected via FACS. RNA was reverse transcribed to cDNA using the High-Capacity cDNA Reverse Transcription Kit (ThermoFisher, 4368814) according to manufacturer instructions with SUPERase-In RNasae Inhibitor (ThermoFisher, AM2694). Mouse Fibrosis RT^2^Profiler PCR Array (Qiagen, PAMM-120ZC, genes in Table A1 below) with PowerUp SYBR Green Master Mix (ThermoFisher, A25777). Cycle run per manufacturer instructions on Applied Biosystems StepOnePlus qPCR system. Results were analyzed with Qiagen supplied data analytics tools available at: https://geneglobe.qiagen.com/us/analyze/. *B2m*, *Gapdh*, *Gusb*, and *Hsp90ab1* were used as reference genes. Data presented as Normalized Fold Regulation over average UI control expression levels.

### Cell culture

24-well plates (Ibidi, 82401) were coated in collagen and laminin at 9 μg/ml and 10 μg/ml, respectively, in PBS for at least 1 hour at 37°C before aspirating and allowed to dry. Freshly sorted FAPs were seeded at a density of 10,000 cells/well, MuSCs were seeded at 5,000 cells/well. FAPs growth medium was DMEM (ThermoFisher, 11965-118), with 20% FBS (Atlanta Biologicals, S11150H), 1% penicillin-streptomycin (PS, ThermoFisher, 15140-122), and 1% GlutaMAX (ThermoFisher, 35050061). Each day in growth media, cells were supplemented with 2.5 ng/ml b-FGF (Sigma, F0291-25UG). MuSC growth medium was Ham’s F10 Nutrient Mix (ThermoFisher, 11550043) with 20% Horse Serum (Atlanta Biologicals, S12150H), 1% GlutaMAX, 1% PS. MuSCs were supplemented with 2.5 ng/ml b-FGF for days 0-3 in culture. FAPs differentiation medium was DMEM, 5% HS (Atlanta Biologicals, S12150H), 1% PS, 1% GlutaMAX. 50% FAPs media by volume was changed every other day. When noted, TGF-β1 (R&D Systems, 7666-MB-005), TIMP1 (Biolegend, 593702), Ultra-LEAF anti-CD29 (Biolegend, 102235), or Ultra-LEAF isotope control (Biolegend, 400940) were added to FAPs differentiation media or directly to MuSC media. MuSCs were allowed to self-differentiate by serum consumption, media was not changed throughout the culture timeline. For transwell experiments, MuSCs were cultured as described with FAPs added in transwell after 3 days in culture. Transwells had 0.4 μm pores (VWR, 29442-129). FAPs were seeded at 10,000 cell/transwell density.

### Partial least squares discriminant analysis and principal component analysis

Partial least squares discriminant (PLSD) computational analysis were done in MATLAB (Mathworks, Natick, MA) using the PLS or function written by Cleiton Nunes (available on the MathWorks File Exchange) according to previously reported protocols (*47*). Briefly, the data were z-scored and then input into the PLS script. D-PLSRs were run for the 32-plex mouse cytokine Luminex assay and the qPCR Fibrosis Array. Orthogonal rotations were applied to maximally separate groups on latent variables in two-dimensions (LV1 and LV2) created by the PLS algorithm. LV loading plots show the mean and standard deviation of each cytokine or gene’s weighting in each latent variable. Standard deviations for loading plots were calculated by iteratively excluding and replacing samples 1000 times and calculating the D-PLSR model each time.

Principal component analysis (PCA) was done using the PCA function in MATLAB. Orthogonal rotations were applied to maximally separate groups on principal components in two-dimensions (PC1 and PC2).

### Uniform manifold approximation and projection (UMAP)

UMAPs were generated as previously reported (*46*). Briefly, UMAP was used here to embed high-dimensional data into a space of two dimensions, and cells are visualized in a scatter plot, where similarity is demonstrated via proximity to other points. Prior to UMAP dimensional reduction, each flow cytometry sample was pre-gated to select for FAPs and then imported into Python 3.7 using fcsparser (https://github.com/eyurtsev/fcsparser) and Pandas 2.5. Each channel except for FSC and SSC was normalized by applying arcsinh/150. A composite UMAP projection that utilized data points from all desired samples was generated using Matplotlib (https://github.com/lmcinnes/umap). Pre-gated cell subsets were overlaid by group onto the UMAP projection to identify the clustered events.

### Spanning-tree progression analysis of density-normalized events (SPADE)

SPADE trees generated as previously described (*46*). Briefly, SPADE was performed through MATLAB and the source code is available at http://pengqiu.gatech.edu/software/SPADE/. MATLAB-based SPADE automatically generates the tree by performing density-dependent down-sampling, agglomerative clustering, linking clusters with a minimum spanning-tree algorithm and up-sampling based on user input. The SPADE tree was generated by exporting uncompensated pre-gated FAPs. The following SPADE parameters were used: Apply compensation matrix in FCS header, Arcsinh transformation with cofactor 150, neighborhood size 5, local density approximation factor 1.5, max allowable cells in pooled downsampled data 50000, target density 20000 cells remaining, and number of desired clusters 100.

### Statistical Analysis

All statistical analyses were done in GraphPad Prism 8. For continuous data with normal distributions and equal variance, students t-tests, one-way ANOVA, or two-way ANOVA were used. Where applicable, a post-hoc test for multiple comparisons was performed, and *p*<0.05 was considered significant. To correct for instances of unequal variance and non-normality of cell frequency data, 2-way ANOVA with Tukey test for multiple comparisons was performed on log-10 or square root transformed data*, p*<0.05 considered significant. Data points are displayed with outlined bars representing the mean, error bars are ± Standard Error of the Mean (SEM).

## Acknowledgments

We thank the core facilities at the Parker H. Petit Institute of Bioengineering and Bioscience at Georgia Institute of Technology and the Atlanta Veterans Affairs Medical Center for the use of shared equipment, services, and expertise.

## Author contributions

SEA, LAH, EAB, YCJ, and NJW designed the research studies, analyzed the research data, wrote the manuscript, and reviewed the manuscript. SEA, LAH, HZ, JMM, TCT, MM, WMH, NHL, JJC, GJ, EG, PC, and SL performed the experiments, analyzed the data, and reviewed the manuscript. GG and LBW contributed to the methodology and reviewed the manuscript.

## Data and materials availability

All data are available in the main text or the supplementary materials.

