## Supplementary Materials for "Aberrant Fibro-Adipogenic Progenitor Subpopulations Drive Volumetric Muscle Loss-Induced Fibrosis"

Shannon E. Anderson *et al.*

**This PDF file includes:**

Figs. S1 to S4

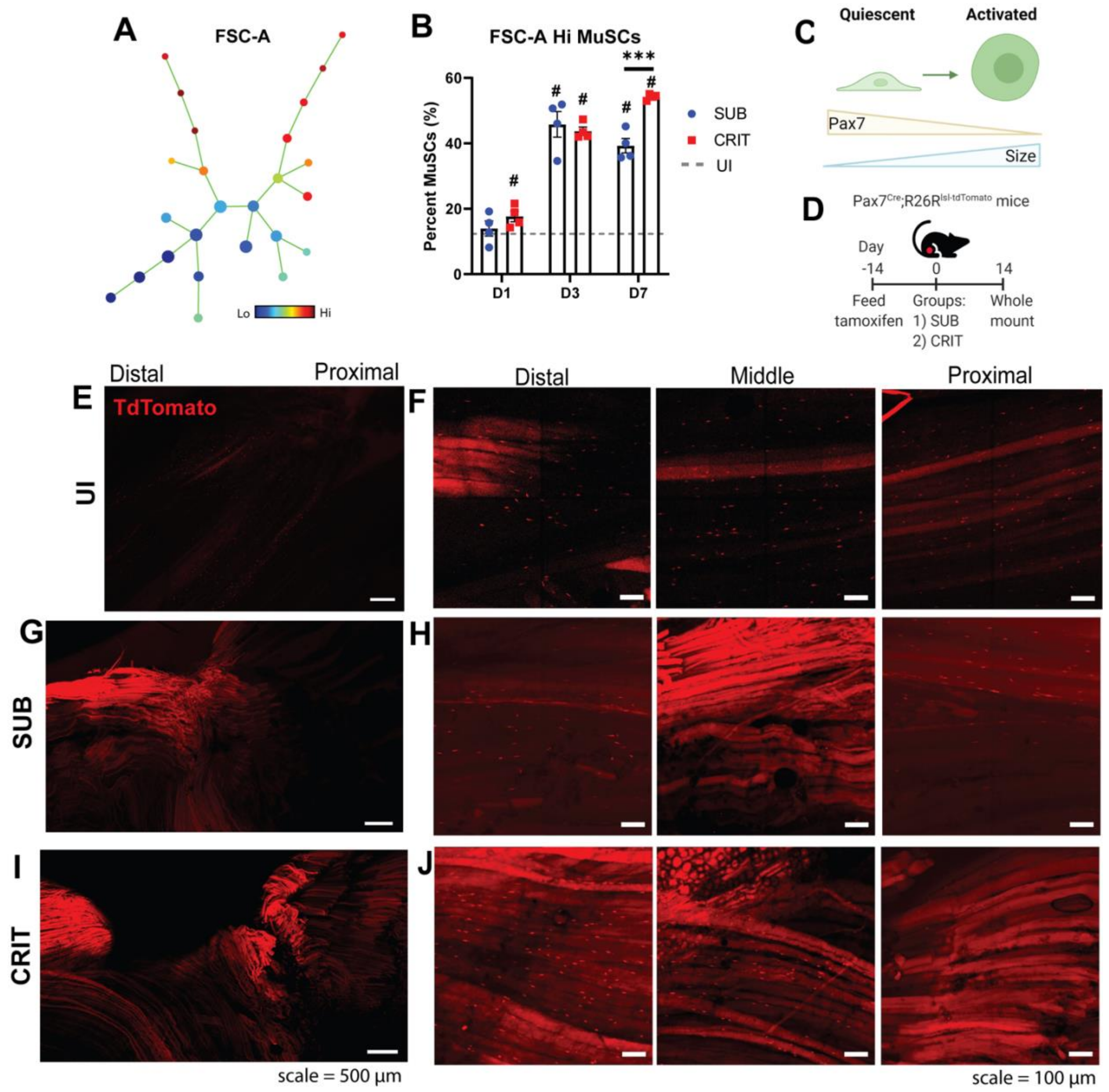


**Fig. S1. Subcritical and critical VML injuries induce MuSC activation and fusion into nearby myofibers**.

A SPADE tree with 25 nodes was created to cluster MuSCs after critical and subcritical injuries over time (**A**) and was used to segment MuSCs into FSC-A high and low groups. The percentage of MuSCs located at FSC-A high nodes was quantified by group (**B**). FSC-A measures cell size, which in the case of MuSCs correlates with activation out of a quiescent state (**C**). Pax7TdT mice were used for whole mount longitudinal imaging (**D**) to track MuSC presence and fusion. Confocal images were taken of each group at different locations along the quadriceps. Sections from uninjured control (UI, **E** and **F**), subcritical injury (SUB, **G** and **H**), critical injury (CRIT, **I** and **J**). Two-way ANOVA was performed using Sidak post-hoc test for multiple comparisons. ***=$p<0.001$, # indicates difference between uninjured (UI) control, $p<0.05$. n=4 animals per experimental group.

**A**

| Fibroblasts 1 | Fibroblasts 2 | Fibroblasts 3 | Tenocytes | FAPs |
|---|---|---|---|---|
| Mest | Gm26917 | Stmn1 | Fmod | Clec3b |
| Ptn | Neat1 | Ube2c | Tnmd | Ly6a |
| Csrp2 | Dst | Cks2 | Scx | Gsn |
| H19 | Kcnq1ot1 | Hmgb2 | Comp | Ly6c1 |
| C1qtnf3 | mt-Atp6 | Birc5 | Ecrg4 | Tnxb |
| Postn | Syne1 | H2afz | Chad | Myoc |
| Mdk | mt-Co3 | Cenpa | Cilp2 | Cd34 |
| Ddah1 | mt-Cytb | Ube2s | Col11a1 | Igfbp5 |
| Col8a1 | Macf1 | Tubb4b | Thbs4 | Lbp |
| Tnc | mt-Co1 | Hmgn2 | Abi3bp | Serping1 |

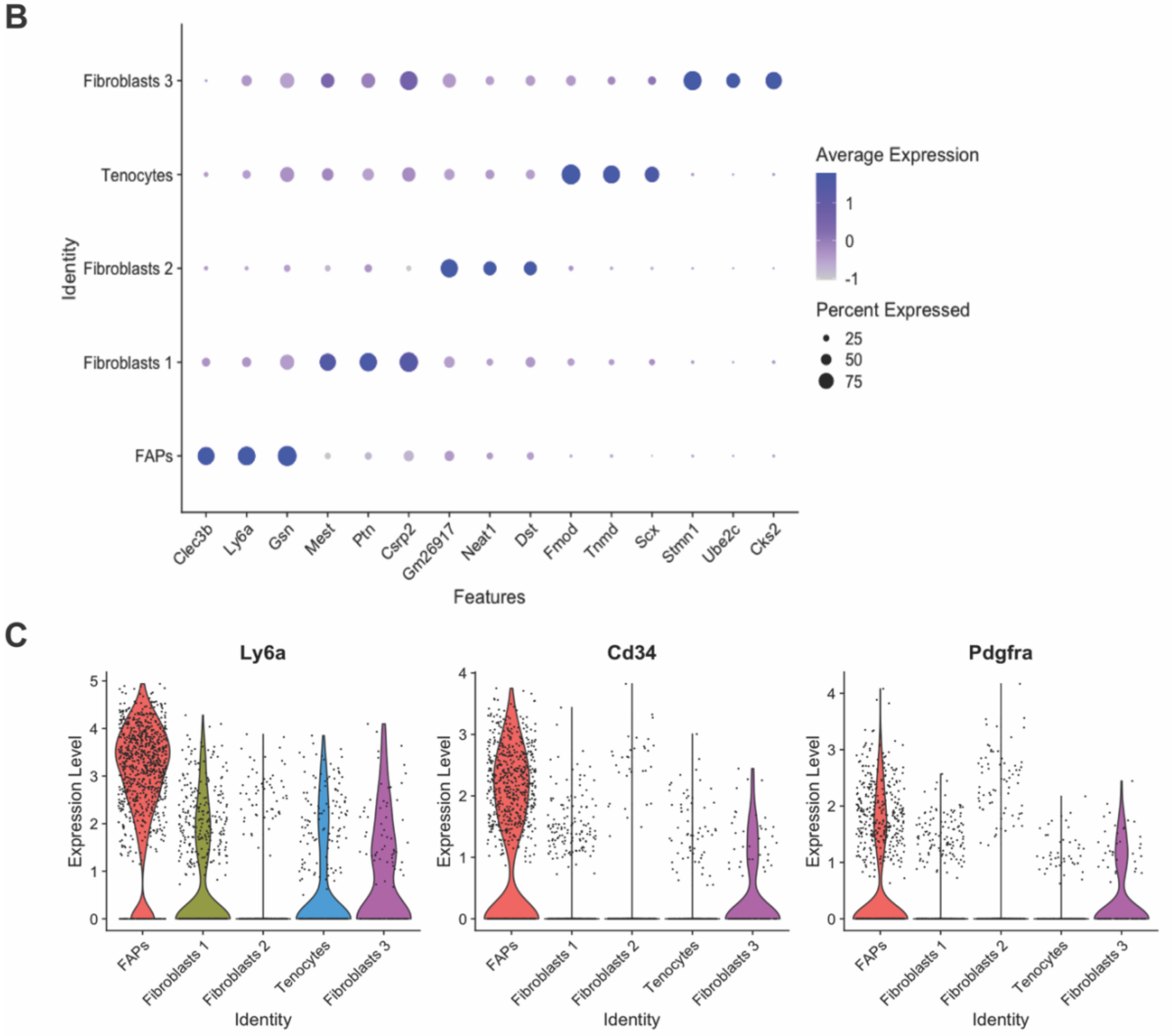


**Fig. S2. Gene expression signatures for identified extracellular matrix cell subsets.**
Table of top 10 genes (**A**) expressed in each of the 5 identified extracellular matrix cell types from quadriceps muscles 7 days post injury or in uninjured controls. Dot plot rendering (**B**) of the top 3 genes expressed per ECM cell type. The size of the dot represents the percentage of cells expressing the indicated gene in a given subcluster. Average gene expression denoted by color scale of gray to purple dot, representing low to high expression, respectively. Violin expression level plots (**C**) for characteristic FAP genes (*Ly6a*, *Cd34*, and *Pdgfra*) for each of the ECM cell types to confirm FAP identification.

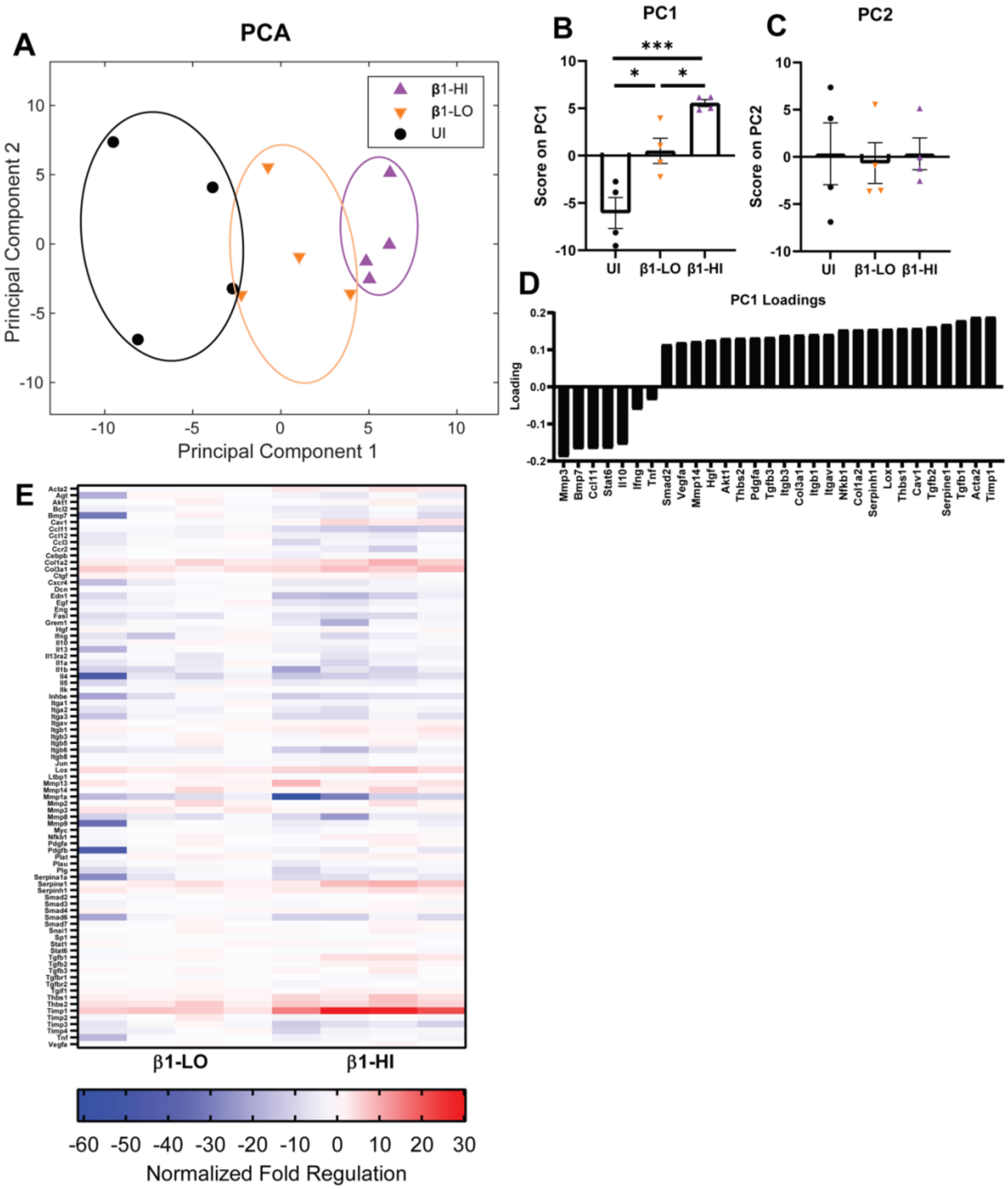


**Fig. S3. Principal component analysis of FAPs fibrosis qPCR array separates UI, β1-LO, and β1-HI FAPs on one principal component.**

Gene expression measured by qPCR from isolated FAPs from uninjured (UI) or injured tissue, with injured FAPs segmented into β1-LO and β1-HI expressing groups was used for PCA and each sample was plotted according to its principal component (**A**). Scores on PC1 (**B**) and PC2 (**C**) were plotted and compared. Differences were seen along PC1, so the loading plot for relative gene contribution to PC1 for select genes is shown (**D**). (**E**) Heatmap showing relative gene expression in β1-LO and β1-HI FAPs. Data quantified as normalized fold regulation relative to uninjured FAPs. One-way ANOVA with Tukey's test for post-hoc, *=$p<0.05$, ***=$p<0.001$. n=4 animals per experimental group.

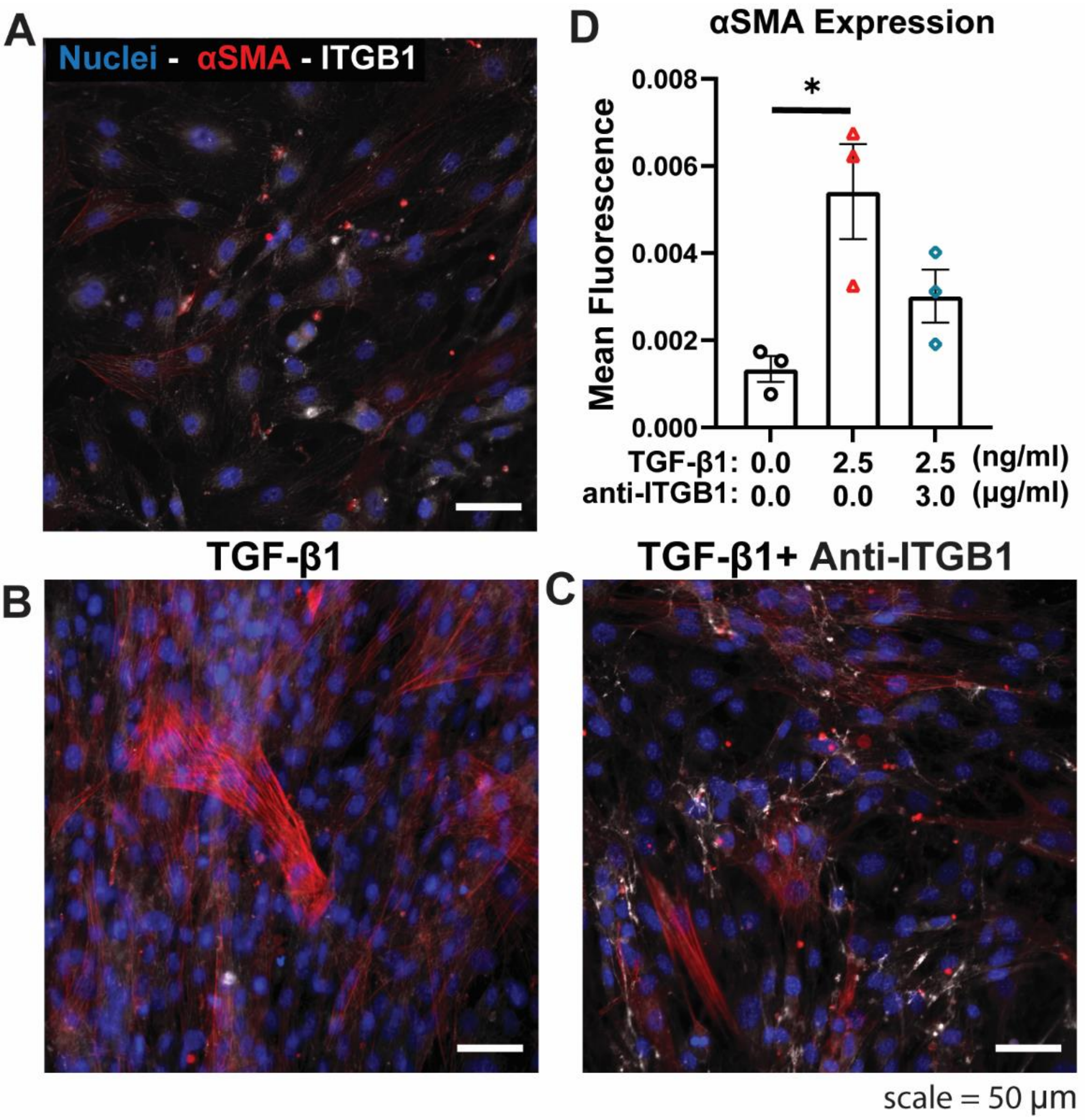


**Fig. S4. Anti-integrin-β1 blocks pro-fibrotic differentiation of FAPs.**

FAPs were isolated, cultured, and differentiated before fixation and IHC staining. Representative images are shown for FAPs with no additives (**A**), TGF-β1 (**B**), or TGF-β1 plus an antibody for Integrin-β1 (Anti-ITGB1, **C**). Staining was for nuclei (blue), αSMA (red), and ITGB1 (white). Mean αSMA fluorescence was quantified (**D**). One-way ANOVA with Tukey's test post-hoc, *=$p<0.05$. n=3 replicates per experimental group.
